# Methods Matter -- Standard Production Platforms for Recombinant AAV Produce Chemically and Functionally Distinct Vectors

**DOI:** 10.1101/640169

**Authors:** Neil G. Rumachik, Stacy A. Malaker, Nicole Poweleit, Lucy H. Maynard, Christopher M. Adams, Ryan D. Leib, Giana Cirolia, Dennis Thomas, Susan Stamnes, Kathleen Holt, Patrick Sinn, Andrew P. May, Nicole K. Paulk

**Author notes:** Corresponding Author: Professor.

## Abstract

Different manufacturing approaches have been used in the production of recombinant adeno-associated virus (rAAV). The two leading approaches are transiently transfected human HEK293 cells and live baculovirus infection of *Sf9* insect cells. Unexplained differences in vector performance have been seen clinically and preclinically. Thus, we performed for the first time a highly controlled comparative production analysis varying only the host cell species but keeping all other rAAV production parameters the same. We demonstrate that host cell species is critical for determining vector potency. Given these key findings, we then sought to deeply characterize differences in rAAVs when produced by these two manufacturing platforms with multiple analytical approaches including: proteomic profiling by mass spectrometry, isoelectric focusing, cryo-EM, denaturation assays, genomic and epigenomic sequencing of packaged genomes, human cytokine profiling, and comparative functional transduction assessments *in vitro* and *in vivo*, including in humanized liver mice. Using these tools we’ve made two major discoveries: 1) rAAV capsids have post-translational modifications (PTMs) including glycosylation, acetylation, phosphorylation, methylation and deamidation, and these PTMs differ between platforms; 2) rAAV genomes are methylated during production, and these methylation marks are also differentially deposited between platforms. In addition, our data also demonstrate that host cell protein impurities differ between platforms and can have their own PTMs including potentially immunogenic N-linked glycans. We show that human-produced rAAVs are more potent than baculovirus-*Sf9* vectors in various cell types *in vitro* (*P* < 0.05-0.0001), in various mouse tissues *in vivo* (*P* < 0.03-0.0001), and in human liver *in vivo* (*P* < 0.005). Collectively, our findings were reproducible across vendors, including commercial manufacturers, academic core facilities, and individual laboratory preparations. These vector differences may have clinical implications for rAAV receptor binding, trafficking, expression kinetics, expression durability, vector immunogenicity as well as cost considerations.

---

Adeno-associated virus (AAV) is a single-stranded DNA virus that is non-pathogenic to humans, exhibits low immunogenicity but high transduction efficiency, and is unable to replicate itself^1^. Recombinant AAV can stably express gene products from either unintegrated episomes^2^ in quiescent tissues, or via integration in actively dividing tissues^3^ when designed with appropriate homology arms. Gene therapies and passive vaccines with rAAV are rapidly gaining attention and investment following the first rAAV therapy approvals in the U.S. market. Nearly 200 rAAV clinical trials for various indications have been performed and numerous investigational new drug applications are in various stages of review at the Food and Drug Administration (FDA) and European Medicines Agency (EMA). The predominant rAAV production platform used for preclinical and clinical studies to date has been transient transfection of adherent human HEK293 cells^4^. However, growth limitations using adherent HEK293 cells have spurred manufacturing innovations to increase yields. A new platform utilizing live baculovirus infection of *Spodoptera frugiperda* (*Sf9*) insect cells produces an average of 7E4 vector genomes (vg)/cell^5-7^. Several recent clinical trials have used vector from this new platform. One difference between their preclinical validation and clinical trials was a transition in manufacturing platform from human HEK293 to baculovirus-*Sf9* when scale-up was needed. This often corresponded to an increased vector dose being administered to achieve relevant expression, and in some cases patients developed severe adverse events directly related to treatment^8,9^. Thus, while poised to revolutionize treatment of rare and common diseases alike, a thorough characterization of differences in rAAV vector safety and potency produced by the two manufacturing platforms was needed.

We characterized differences in vector lot composition at both the genomic and proteomic levels. This was largely motivated by the fact that human and insect cells have different capacities to produce protein post-translational modifications (PTMs)^10^. Protein folding and PTMs can influence therapeutically administered proteins, including altering stability, targeting/trafficking, functional activity, and immunogenicity, all of which could affect rAAV potency. Additionally, a concern in producing recombinant proteins for humans within insect cells is potential immunotoxicity. Humans can have acute allergenic reactions to non-mammalian N-glycans, as well as any N-glycan with an ɑ1,3-fucose or β1,2-xylose linkage on the innermost N-acetyl glucosamine (GlcNAc)^11^, both of which are modifications found on insect glycoproteins. Thus, certain insect glycoproteins in baculovirus-*Sf9* produced vector lots could pose potential risks.

Impurities of any kind are a known concern for all production methods. For example, the EMA assessment report for the rAAV Glybera™ found the final vector lot impurities to be ‘unacceptably high’^12^. Collectively, we therefore hypothesized that rAAV vector--both capsids and packaged genomes--produced in different manufacturing platforms may be compositionally diverse. Using an array of chemical, molecular, structural, bioinformatic, and functional assessment approaches, we characterized and report here the differences and similarities in rAAV vector produced by these two leading manufacturing platforms.

## Results

### The vector PTM landscape differs in rAAV vectors made using the human and baculovirus-Sf9 production platforms

We sought to determine whether rAAV capsids were post-translationally modified, and whether capsid PTMs or any host cell protein (HCP) process impurities differed between production methods and lots. We used deep proteomic profiling and liquid chromatography-tandem mass spectrometry (LC-MS/MS). To eliminate variables between human and baculovirus-*Sf9* platforms for comparative high-resolution MS/MS analysis, the following safeguards were implemented: a) all vector productions were carried out at the same time in one facility using identical equipment by the same individuals; b) vector lots were harvested, purified and underwent quality control together using the same assays; c) vector aliquots were frozen down identically and simultaneously; and d) aliquots were thawed together and prepared for analysis from lots that had been frozen for the same amount of time to eliminate the effects of time spent frozen on potential PTM retention. One additional parameter that was important to control was the input DNA and backbone vector sequence itself. Recombinant AAV transfer vectors are encoded on plasmids for transfection in the human manufacturing platform and within a live baculovirus in the insect platform. To overcome this difference, we designed a custom transfer vector plasmid that has the necessary backbone components to be employed in both systems (**Figure S1**). Thus, the only differences remaining were those standard to each method: human HEK293 cells were grown in adherent cultures and transiently transfected with three production plasmids, while insect *Sf9* cells were grown in suspension cultures infected with two second generation^5^ recombinant baculoviruses needed to produce rAAV.

A set of rAAV8 productions was carried out and subdivided into four preparations for simultaneous purification: lots purified from cell lysates and media supernatant from each platform. LC-MS/MS analysis resulted in rAAV8 capsid protein coverage of 98.8% (human) and 97.3% (baculovirus-*Sf9*). Capsid modifications observed included O-linked glycans, N-terminal acetylation to start methionine, lysine/arginine acetylation, tyrosine/serine/threonine phosphorylation, lysine/arginine methylation, and aspartate deamidation (**Figure S2a-f**). PTMs were distributed along the entire polypeptide, and more PTMs were observed on baculo-rAAV8 capsids compared to those produced in human cells (**Figure 1a-e**). In addition, externally facing PTMs were more common than those in the capsid lumen in both platforms. However, the first 220 amino acids in the N-terminus of rAAV8 are disordered; thus, we lack structural data to classify the orientation of PTMs in that region. Limited data suggests that these amino acids are located within the capsid^13,14^. Interestingly, several PTMs within this disordered and presumed lumenal region were shared between the human and baculo-rAAV8 lots: N-terminal acetylation on the initial start methionine, and phosphorylation on serines 149 and 153. These shared PTMs may play a conserved role in capsid-genome interactions or in intracellular trafficking following phospholipase A2 domain extrusion once in the endosome^15^. We confirmed the presence of all 16 recently reported deamidations which were observed on human cell-lysate purified rAAV8^16^ on both human and baculo-rAAV8 cell lysate and media-purified preparations.

**Figure 1.**
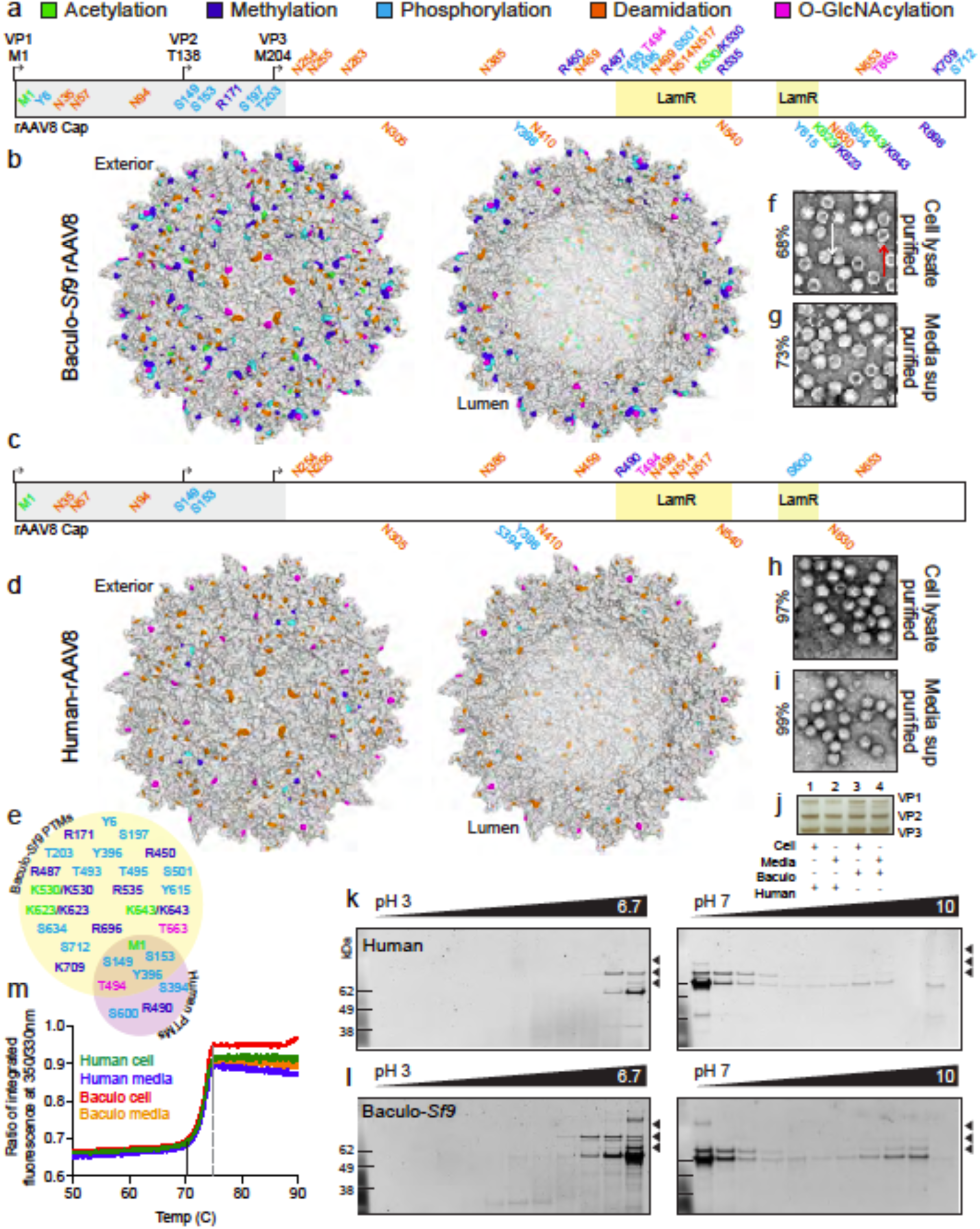
rAAV capsids manufactured with the human and baculovirus-*Sf9* platforms are post-translationally modified and exhibit differential PTM profiles. **(a)** PTM identities and residue positions along the length of the rAAV8 polypeptide from N to C-terminus in baculo-*Sf9* vector. PTMs colored by type (acetylation = green, methylation = blue, phosphorylation = cyan, deamidation = orange, O-GlcNAcylation = magenta). Residues above the sequence are externally facing on the capsid. Residues below are lumenal or buried. Residues within the grey box from 1-220 represent the disordered region of AAV8 yet to be crystallized. The two regions for LamR binding (491-547 and 593-623) are highlighted in yellow boxes. **(b)** Cumulative capsid PTMs observed from all baculo-*Sf9* rAAV8 lots, purified from both cell lysates and media. Same color code as in (a). **(c)** Same as (a) but with human-produced rAAV8. **(d)** Same as (b) but with human rAAV8. **(e)** Shared and unique capsid PTMs for rAAV8 produced in the baculo-*Sf9* (yellow) and human platforms (purple). Same color code as in (a). Excluded are deamidation degradation marks which are universal. **(f)** Negative staining and TEM imaging of baculo-*Sf9* rAAV8 cell-purified vector. White arrow = full capsid, red arrow = empty capsid, for reference for panels f-i (percent full capsids noted on left). **(g)** Same as (f) but media purified vector. **(h)** Same as (f) but with human rAAV8 cell-purified vector. **(i)** Same as (h) but with media-purified vector. **(j)** Silver stain of capsid VP species present in vector lots from panels f-i. **(k)** 2D gel images from human-produced rAAV8 from pH 3-10. VP1 (87 kDa), VP2 (72 kDa), and VP3 (62 kDa) bands = black arrowheads. **(l)** 2D gel images from baculo-*Sf9* produced rAAV8. **(m)** Thermal capsid melt curves for rAAV8 vectors shown from 50-90°C, full melt curves from 30-95°C are in Fig S7a. Tm initiation = dashed black line; final Tm = dashed grey line.

For the remaining residues 220-738 on rAAV8 for which robust structural data exists^17^, those residues which are externally facing may play roles in mediating blood clearance, antibody binding, cellular tropism, receptor binding, etc. We detected numerous PTMs within known external functional regions (**Figure 1a-e**). For example, within the laminin receptor (LamR) binding region (residues 491-547 and 593-623)^18^, we observed one shared and numerous differential PTMs between human and baculo-produced vectors. The one conserved modification was an O-GlcNAc on T494 within the LamR binding domain. However, baculo-rAAV8 had five different external modifications, compared to the two in human-rAAV8 in this same domain. This included K530, which we detected to be capable of both acetylation and methylation on different capsids within the same vector lot. Interestingly, the C-terminal end of the human-produced vectors lacked observable PTMs, both internally and externally, while the baculo-produced rAAV8 vectors were often modified in this region, including an additional external O-GlyNAc on T663. No PTMs were detected in the known ADK8 neutralizing epitope region (586-591)^19^. Additionally, there are 8 lysines on the mapped structure of AAV8 VP3 predicted to be capable of ubiquitination^20^: K259, K333, K510, K530, K547, K569, K668 and K709. In human rAAV8 productions we observed no PTMs on or near any of these residues, while in baculovirus-*Sf9* productions we observed PTMs on two ubiquitinable lysines (K530 was methylated and acetylated; K709 was methylated), thus potentially blocking them from future ubiquitination.

Negative staining and transmission electron microscopy (TEM) confirmed that, despite identical purification, human-rAAV had more full capsids, regardless of whether the rAAVs were purified from cell lysates or media supernatant (**Figure 1f-i**). We observed normal 1:1:10^21^ ratios of VP1/2/3 by silver stain (**Figure 1j**) for each vector lot, however the baculovirus-*Sf9* vectors displayed additional truncated VPs, as seen by others previously^22^. HCP impurities were present in all vectors, regardless of manufacturing platform, but were different between platforms (**Table S1**). Of concern, we observed *Sf9* insect HCP impurities with N-linked glycans (**Table 1, Figure S2g,h**). Given that humans can have acute allergenic reactions to non-mammalian N-glycans^11^, the insect N-glycans found in baculovirus-*Sf9* produced vector could pose potential risks. In addition, we performed cytokine profiling in primary human fibroblasts following transduction with each of the four vector lots by Luminex assay and demonstrated that responses were more similar for human-produced rAAV8 vectors than for baculo-*Sf9* vectors (**Figure S3**). Specifications on the viral lots are outlined in **Table S2**.

**Table 1.**
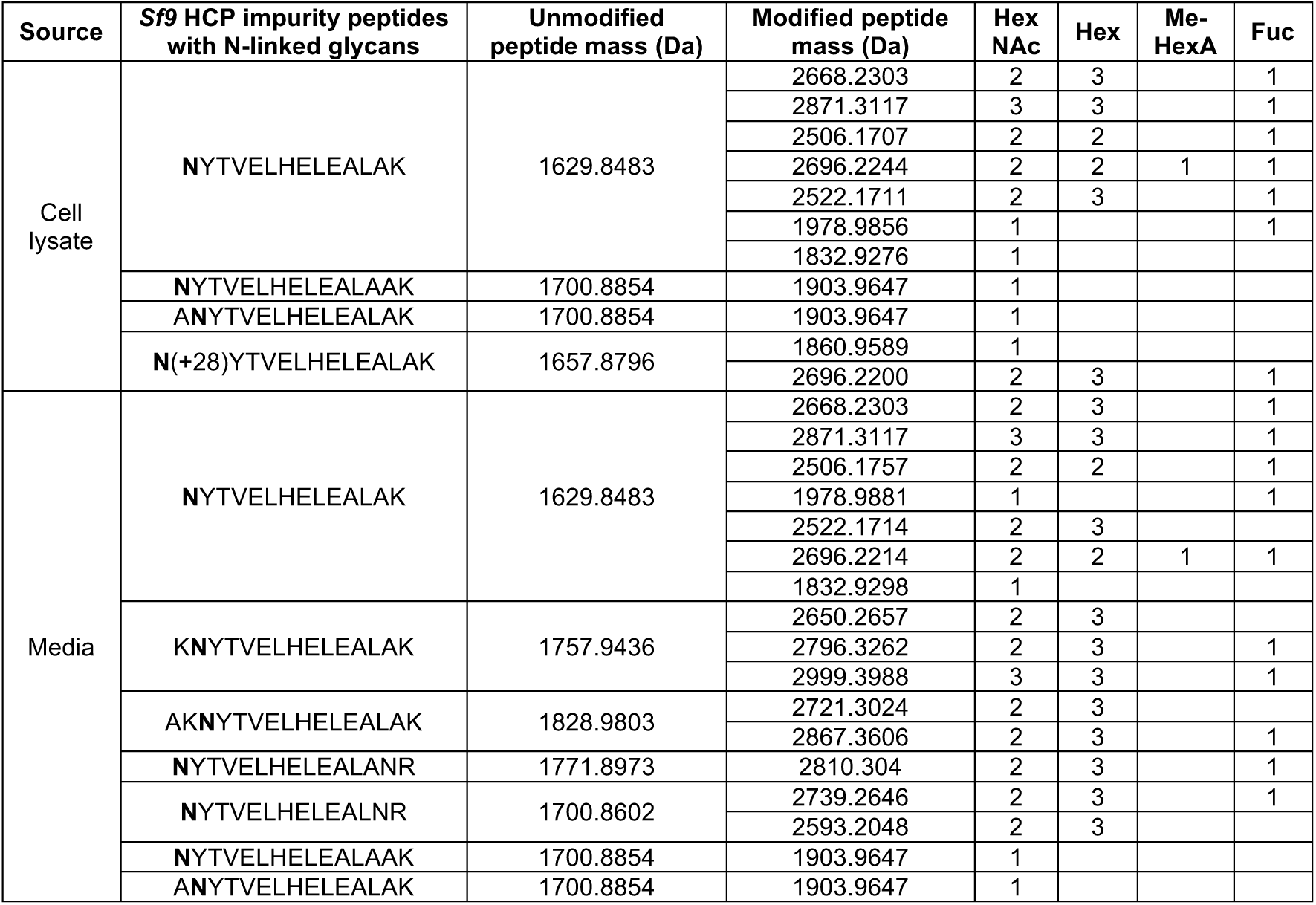
Host cell protein impurities with N-linked glycans from baculovirus-*Sf9* productions of rAAV8. N-linked glycans (modified residue = **bold**) identified on the common *Sf9* HCP impurity ferritin identified by LC-MS/MS for rAAV8 from both cell lysate and media-purified vectors. Excluded from the list are common process contaminants that occur in routine sample preparation (human keratin, trypsin, etc.), impurities with mutations/aberrations such that they didn’t map to known proteins by BLASTp search, and any modification which could not be site-localized. Unmodified and modified peptide masses are shown, as well as the number of glycan moieties each modified mass represents. HexNAc = N-acetylhexoseamine; Hex = hexose; Me-HexA = methylated hexuronic acid; Fuc = fucose; Da = dalton. No HCP impurity peptides in the human rAAV8 preparations were observed with N-linked glycans.

To determine whether the baculovirus-*Sf9* rAAV results would be different with first generation^5^ baculoviral constructs—as were used in the rAAV1-LPL trial for Glybera^23^—we prepared another four lots of rAAV8, only now using the original baculoviral support constructs for the baculovirus-*Sf9* productions. Here again, results with first generation constructs were consistent with those reported for the former replicates using second generation constructs (**Figure S4, Table S3**). Host cell ferritin contamination of suspension-grown rAAV preparations is known^24^, and was seen in these baculovirus-*Sf9* lots.

### 2D gel electrophoresis independently confirmed vector differences

To validate differences observed at the proteome level from LC-MS/MS, we performed 2D gel electrophoresis (in-solution isoelectric focusing followed by SDS-PAGE) on four paired rAAV8 vector samples from Figure 1. Briefly, samples were loaded into wells above a pre-cast pH gradient gel strip and run in the presence of current such that proteins migrated through the gel to the well where the isoelectric point (pI) matched the well pH. After migrating, samples in each well were removed and run in separate lanes on a standard SDS-PAGE gel to separate by size in the second dimension (**Figure S5**). Chemical differences affecting pI in capsid proteins manifest as lateral banding, and we observed VP1/2/3 proteins that migrated beyond the well corresponding to the natural pI of rAAV8 (pI 6.3)^25^ (**Figure 1k,l**).

These bands reflect differences in PTMs or other unidentified modifications. In different conditions (pH/temperature), rAAV can undergo conformational changes due to phospholipase A_2_ catalytic domain flipping within VP1^25^. However, this would not result in the observed lateral banding, as conformational changes do not significantly alter net charge, and thus pI, of a protein^26^. Additionally, lateral banding was observed predominantly on VP3, which lacks the phospholipase domain. Of note, VP3 bands had substantially greater lateral banding than VP1 or VP2, possibly due to its higher abundance. To rule out the potential influence of host cell proteases being responsible for the banding patterns seen by 2D gel electrophoresis, an independent control was run wherein rAAV8 samples were treated with and without protease inhibitors and run as before to ensure the banding patterns remained unchanged. No difference was seen between treatments (**Figure S6a**) confirming the previous findings that banding patterns were not due to contaminating proteases. To determine if lateral banding resulted from rAAV cleavage products or substantial loss of PTMs, immunoblot analysis of the 2D gels was performed with an anti-AAV VP1/2/3 antibody^27^. Immunoblots confirmed that lateral banding was rAAV capsid proteins and that vertical banding was likely attributable to HCPs (**Figure S6b,c**). To determine whether differences imparted by capsid PTMs could influence capsid stability, thermal melt curves were performed and no differences were found between baculovirus-*Sf9* and human-produced rAAV8 (**Figures 1m, S7a**). All displayed standard rAAV8 melt curves initiating at 70°C and completing at 75°C, similar to rAAV8 vector lots from other academic and commercial providers (**Figure S7b-d**) for each platform. Of note, every rAAV serotype, regardless of method of production, has a unique capsid Tm that can vary from as low as ∼72°C for rAAV2 to as high as ∼91°C for rAAV5, with other common serotypes falling in between (**Figure S7b-d, Table S4**).

### Capsid alterations and HCP impurities are consistent across serotypes

To understand whether the former results with rAAV8 were serotype-specific or inherent to each platform, an identical set of rAAV1 preparations was produced and subdivided into four separate groups for simultaneous purification as was done for rAAV8. LC-MS/MS results, with an average of 87.9% (human) and 83.4% (baculovirus-*Sf9*) capsid coverage, showed that rAAV1 capsids are also post-translationally modified, and that the types and frequencies of each again differed between the two platforms (**Figure S8a-e**). Four rAAV1 PTMs within the disordered N-terminal region of the capsid were shared between the human and baculo-rAAV1 vectors: N-terminal acetylation on the initial start methionine, methylation of lysine 61, and phosphorylation on serines 149 and 153. For the remaining residues 217-736 on rAAV1, we detected numerous PTMs within known antigenic motifs that react with known neutralizing antibodies against rAAV1 (456-AQNK-459), (492-TKTDNNNS-499), and (588-STDPATGDVH-597)^28^. One PTM was detected within the 4E4 neutralizing epitope residue (456-459 and 492-498)^29,30^, and several additional PTMs were detected near the 5H7 neutralizing epitopes (494, 496-499, 582, 583, 588-591, 593-595, 597), as well as the ADK1a neutralizing epitope (500). Additionally, there are 11 lysines on the mapped structure of rAAV1 VP3 predicted to be capable of ubiquitination: K258, K459, K491, K493, K508, K528, K533, K545, K567, K666 and K707. In both human baculovirus-*Sf9* rAAV1 productions we observed PTMs on two ubiquitinable lysines where K459 and K528 were methylated. Here again, HCP impurities were present and different between platforms (**Table S5**), and *Sf9* insect HCP impurities were detected with N-linked glycans (**Table S6**). Specifications on the viral lots are outlined in **Table S7**. Negative staining and TEM imaging confirmed that, despite identical purification procedures, human-produced rAAV1 had fewer HCP impurities and reproducibly full capsids compared to baculovirus-*Sf9* vector preparations (**Figure S8f-i**). We observed normal VP1/2/3 ratios by silver stain (**Figure S8j**), and again did not detect significant differences in thermal capsid stability (**Figures S8k, S7e**).

### rAAV capsids are structurally similar regardless of production method

Having demonstrated differences in composition at both the capsid PTM and vector lot impurity levels when produced by different production platforms, we next wanted to assess whether global capsid structures differed by transmission electron cryomicroscopy (cryo-EM). To fairly assess potential differences, we first needed comparable structures from identically purified vector lots subjected to cryo-EM using the same methodology. Thus, we generated four new cryo-EM structures of full and empty rAAV8 capsids from each manufacturing platform (**Table S8**). Despite differential PTM deposition, no significant differences were observed between the averaged structures at the resolutions achieved (3.3-3.6Å) (**Figures 2, S9**). The 4 new capsid structures have been deposited in both PDB and EMDB: human full AAV8 (PDB-6PWA; EMD-20502); human empty AAV8 (PDB-6U20; EMD-20615); baculovirus-*Sf9* full AAV8 (PDB-6U2V; EMD-20626); baculovirus-*Sf9* empty AAV8 (PDB-6UBM; EMD-20710).

**Figure 2.**
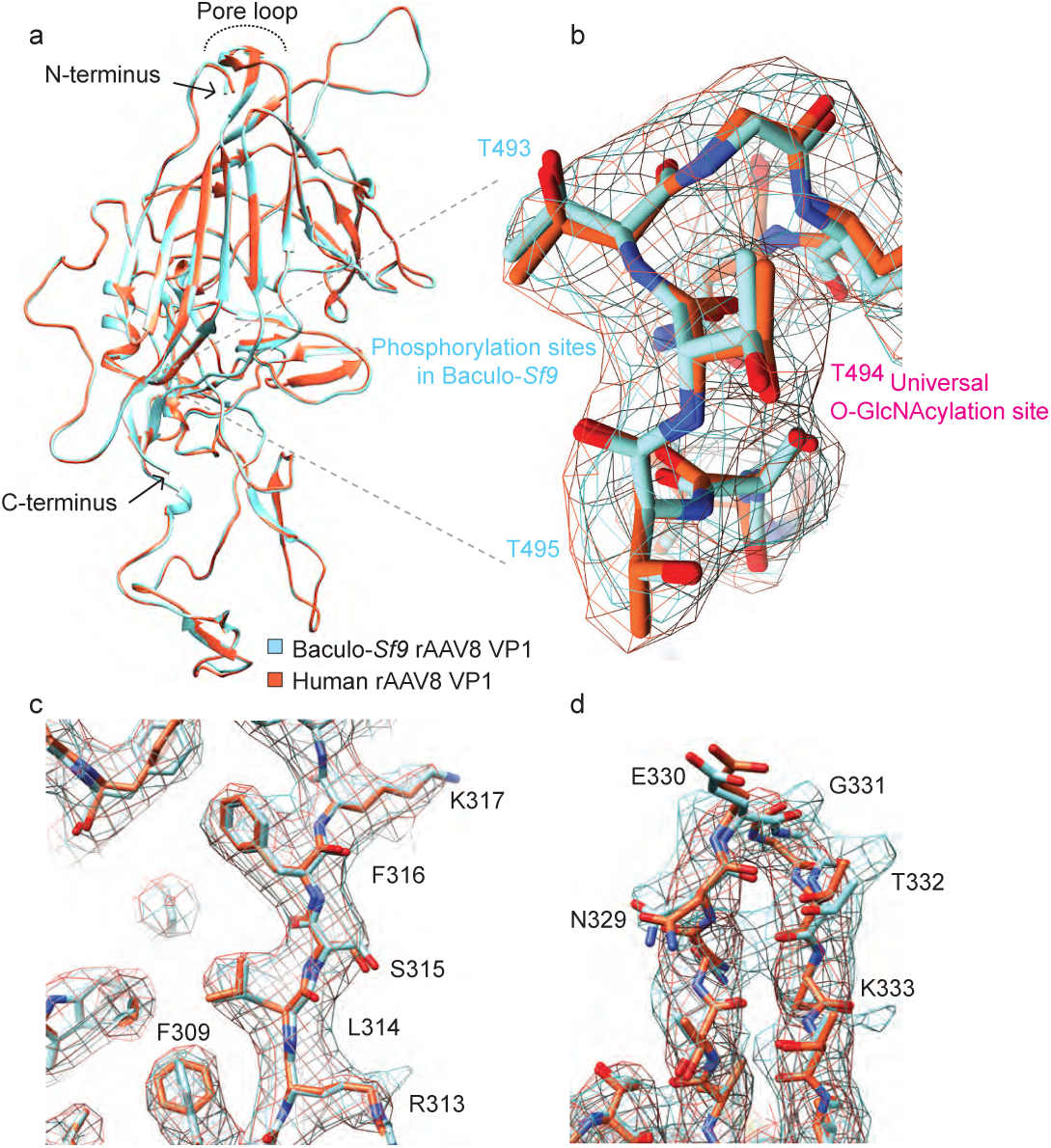
Human and baculovirus-*Sf9* production platforms produce rAAV capsids with similar structures by cryo-EM. **(a)** Overlay of an individual VP1 chain from human (red-orange) and baculo-*Sf9* (cyan) full rAAV8 capsids highlighting an absence of macro-level differences. Key structural landmarks are shown. **(b)** Magnified overlay of the LamR binding region from a human (red-orange electron density caging) and baculo-*Sf9* (cyan) full rAAV8 capsid highlighting residues capable of post-translational modification as determined by LC-MS/MS. **(c)** Magnified overlay depicting potential side-chain level structural differences in phenyl ring orientations between human and baculo-*Sf9* rAAV8. **(d)** Magnified overlay of a single human and baculo-*Sf9* rAAV8 capsid cylinder loop depicting minor potential side-chain level differences.

### Common rAAV serotypes have capsid PTMs and HCP impurities

After having shown that standard rAAV production methods produce capsids decorated with PTMs, we wanted to determine whether this was specific to the two serotypes tested or a generalizable phenomenon. To assess the presence and/or absence of capsid PTMs across serotypes, we sourced a panel of common rAAV serotypes 1-8, produced in either the human or baculovirus-*Sf9* platforms (**Table S9**) from a single manufacturer to minimize potential production variation, and performed LC-MS/MS. We observed that all rAAV capsids, regardless of serotype or manufacturing platform, possessed capsid PTMs (**Table S10**). Additionally, all vector lots had HCP impurities (**Table S11**), and some HCPs were detected with N-glycosylation (**Table S12**).

### Capsid PTM deposition and HCP impurities are universally seen across manufacturing platforms, purification methods, facilities and scale

Up to this point, vector lots from a single manufacturing facility in highly controlled production and purification settings were assessed. We next wanted to determine whether these findings were generalizable across different production batches and manufacturing sites. To examine this, we sourced rAAV vector lots from an array of production facilities spanning private industry, leading academic core facilities, government consortia and individual academic laboratories (**Table S13**). Here the intent was not to rigorously control the production and purification parameters, but rather to determine the extent of modifications and impurities with samples obtained from vendors supplying vectors to scientists working at every stage in the R&D pipeline. LC-MS/MS demonstrated that capsid PTMs and HCP impurities were universal (**Tables S14-15**), regardless of the scale of manufacturing, purification method used, manufacturing facility, or production platform. Cumulative comparative HCP impurity analyses on all vector lots tested to date using gene ontology analysis highlighted trends in HCP impurity types (**Figure S10** and **Tables S16-18**). Interestingly, the most common human impurities were related to nucleic acid and protein binding for RNA processing, the most common insect impurities were related to endopeptidase activity and proteolysis, while baculoviral impurities were either structural in nature or pertaining to viral escape.

Lastly, despite the presence of putative endoplasmic reticulum (ER) signal peptides^31^ at the N-terminus of VP1 on all common rAAV serotypes (**Table S19**), our LC-MS/MS data show that rAAVs are often N-terminally acetylated, a PTM known to block ER translocation of polypeptides in eukaryotes^32^. The potential ER-exclusion is consistent with the glycosylation patterns seen on rAAV capsids which have been observed to have simple O-GlcNAc modifications (deposited in the nucleus/cytoplasm), but lack N-glycans or complex O-glycans (deposited in the ER/Golgi).

### rAAV genomes are differentially methylated during production

Given the numerous unexpected differences in rAAV vectors at the proteomic level, we hypothesized whether similar differences may also be occurring at the epigenomic level. We were particularly interested in quantifying genome methylation, given it’s known role in gene regulation, which could impact vector potency. However, similar to our discovery of capsid PTMs presented earlier, no literature existed on whether rAAV genomes could be methylated during packaging. Here we sought to determine whether packaged rAAV genomes were capable of being epigenetically methylated, and if so, whether this differed between production platforms. We used whole genome bisulfite sequencing (WGBS) to generate genome-wide methylation maps of the rAAV genome at single-base resolution. ssDNA for WGBS was isolated from rAAV vectors from each production platform and unmethylated bacteriophage lambda DNA was spiked into each rAAV sample prior to bisulfite conversion. Sequencing generated 17.3 million raw reads that were subsequently mapped to the rAAV transfer vector and reference genomes. We discovered that rAAV genomes are indeed methylated and that the cytosine methylation was distributed along the entire genome (**Figure 3**). Overall methylation levels were low (∼1%) and similar between the two platforms. However, three sites in particular showed statistically significant differential methylation between vectors from the two platforms: repressive promoter methylation at C:1208 and polyA methylation at C:3779 was significantly higher in baculo-rAAV; while activating intragenic methylation at C:2970 was significantly higher in human-rAAV.

**Figure 3.**
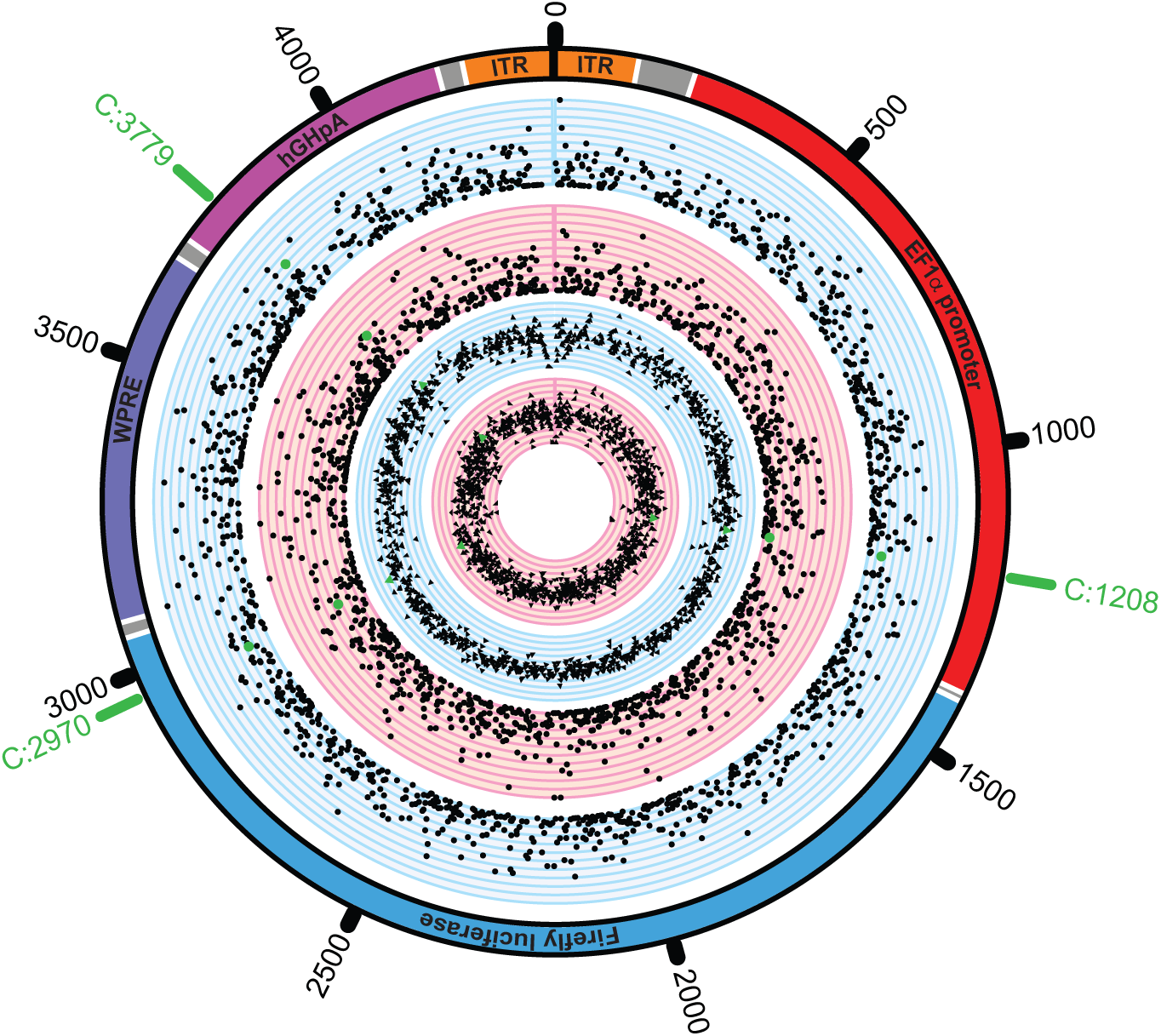
Packaged rAAV genomes are methylated during production and differentially methylated based on manufacturing platform. The five tracks of the circos plot from the outside in represent the following. First, a schematic of the ssAAV8 genome sequenced with key features labeled (inverted terminal repeat (ITR) = orange; intervening sequences = grey; EF1α promoter = red; Firefly luciferase expression cassette = blue; woodchuck hepatitis virus posttranscriptional regulatory element (WPRE) = plum; human growth hormone polyadenylation signal (hGHpA) = magenta). Second, in blue, the methylation ratio for all detected sites for baculovirus-*Sf9* produced rAAV8; the plotting represents a minimum of 0 and a maximum of 1 in 1/10 increments. Third, in red, the methylation ratio for all detected sites for human-produced rAAV, scale same as previous. Fourth, in blue, the total number of reads at detected sites for baculovirus-*Sf9* produced rAAV8; in log2 scale, from a minimum of 0 (total reads of 1) to a maximum of 11 (total reads of 2^11), in 1/10 increments. Fifth, in red, the total number of reads at detected sites for human-produced rAAV, scale same as previous. The three sites with statistically significant differential methylation are highlighted in green (C:1208, C:2970, C:3779).

### Human-produced rAAV exhibits significantly greater potency in vitro

Having established that capsid PTMs, HCP impurities and epigenomic methylation were key differences in rAAV vector lots, we next sought to quantify vector potency differences between manufacturing platforms. We setup comparative expression experiments in a panel of cell lines spanning different species, cell types, differentiation states, and immortalization statuses. Immortalized human HEK293T and Huh7 cells, primary human fetal fibroblasts, primary human iPSCs, and immortalized mouse C2C12 myoblasts were transduced with rAAV1 Firefly Luciferase (FLuc) vectors to measure potential expression differences. FLuc assays confirmed significant potency differences in all cases, regardless of species or cell type, with human-produced rAAV vectors outperforming insect-produced rAAV vectors in all cases (*P <* 0.05-0.0001) (**Figure 4a**).

**Figure 4.**
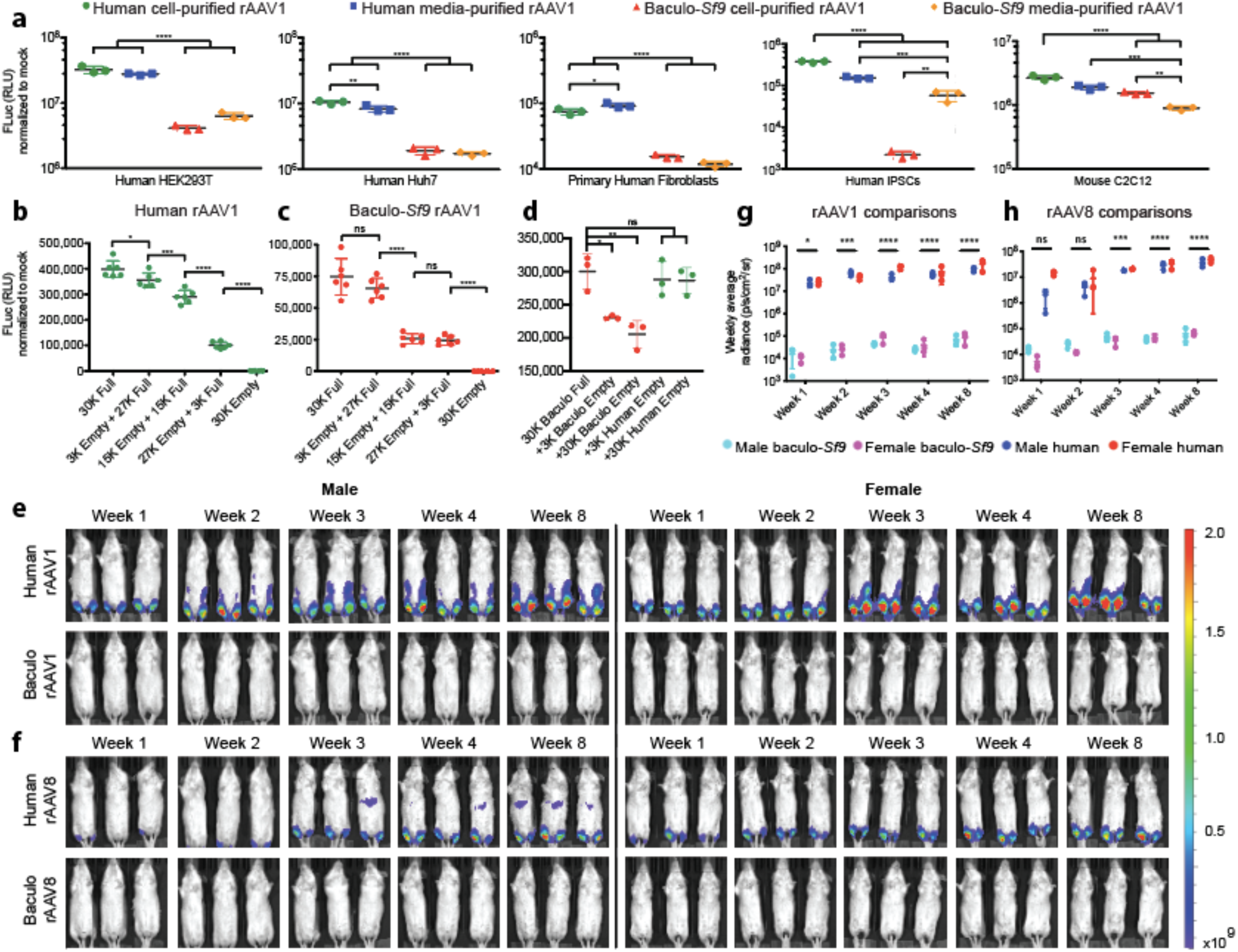
rAAV produced with the human platform is more potent *in vitro* and in skeletal muscle *in vivo* following intramuscular administration. **(a)** *In vitro* functional transduction assays in immortalized human HEK293T and Huh7 cells, primary human fibroblasts, primary human iPSCs, and immortalized mouse C2C12 myoblasts transduced with ssAAV1-EF1α-FLuc. Human-produced rAAV1 was significantly more potent than similar baculovirus-*Sf9* vector in all cases. **(b)** HEK293T cells were transduced with varying ratios of full:empty human-produced ssAAV1-EF1α-FLuc with the total capsid content kept constant at MOI 30K, while the ratio of full:empty varied. FLuc assays were performed on cell lysates 3 days post-transduction and normalized to mock-transduced wells. Each green dot represents one replicate; mean +/- SD shown. **(c)** Same as (b) except using baculovirus-*Sf9* rAAV1. **(d)** To further assess the impact of insect HCP impurities, HEK293T cells were transduced with a fixed 30K MOI of full baculo-produced ssAAV1-EF1α-FLuc and spiked with an additional 10% or 100% of empty baculo-produced or human-produced vector. FLuc assays were performed on cell lysates 3 days post-transduction and normalized to mock-transduced wells. Each dot represents one replicate; mean +/- SD shown. **(e)** *In vivo* time course functional transduction assays comparing human and baculovirus-*Sf9* ssAAV1-EF1α-FLuc after IM administration (5E10 vg/mouse) in age-matched siblings. Mean radiance (p/s/cm^2^/sr) displayed with all mice imaged on their ventral side on the same shared scale. **(f)** Same as (e) but with ssAAV8-EF1α-FLuc; same shared scale as (e). **(g)** Quantification of rAAV1 FLuc radiance from (e). Each symbol = mean signal (+/- SD) from 3 mice. **(h)** Same as (g) but with rAAV8. * *P* ≤ 0.05, ** *P* ≤ 0.01, *** *P* ≤ 0.001, **** *P* ≤ 0.0001.

To validate that expression differences were not due to differential packaging efficiency, we developed a next-generation sequencing (NGS) protocol called Fast-Seq^33^ to sequence packaged rAAV genomes, including ITRs. Fast-Seq validated that all vectors used in the potency assessments had no difference in the packaged genomic sequence between any vector lot by either production method (**Figure S11** and **Tables S20-21**). These findings substantiate that observed expression differences were not due to differences in genome sequence or integrity.

To assess whether host cell protein impurities and/or empty capsids were poisoning the potency of the baculovirus-*Sf9* preparations, we setup an *in vitro* experiment with rAAV1-FLuc wherein the total capsid MOI was kept constant and standard genome-containing capsids were mixed with purified lots containing only empty capsids and their associated HCPs at different ratios (100:0, 90:10, 50:50, 10:90, 0:100) to determine whether the signal drop-off would correlate with the percent of empty vector and/or HCP impurities. Human-rAAV1 expression dropped less quickly (**Figure 4b**) than baculo-produced rAAV1 (**Figure 4c**) when >10% of the total vector material came from empty vector/HCP impurities. To further probe the influence of insect/baculoviral impurities, we setup a similar experiment wherein the capsid MOI was not fixed but we spiked empty capsids and HCPs on top of the existing 30K full baculoviral capsids and again similarly assessed transduction. If insect/baculoviral HCP impurities and/or empty capsids were not influencing potency, then the spike-ins should have no effect. However, as is seen with 10% or 100% baculoviral empty spike-ins, we observed a significant loss of expression from the full capsids (**Figure 4d**). Yet similar spike-ins with human empties had no effect, and also demonstrated that empty capsids where not merely outcompeting full capsids for available receptors. Together, these experiments support the hypothesis that the presence of insect/baculoviral impurities and/or empty capsids in baculovirus-*Sf9* preparations is influencing the potency of these vector preparations.

### Human-rAAV has significantly greater muscle transduction in vivo

To assess potency differences *in vivo*, we setup a transduction comparison with the following parameters: a) age-matched mice to prevent confounding effects of age-related transduction; b) mice of each sex were treated; c) the same ssAAV-CMV-FLuc-SV40pA transfer vector was used to facilitate comparisons between production methods and manufacturers; d) vectors were sourced from two popular manufacturers for each production platform (viral lot specifications in **Table S22**); e) all mice were injected intramuscularly (IM) by the same person on the same day and housed in their own cage per treatment group to prevent cross-contamination by vector shedding; f) we used the same serotype (rAAV1) and method of injection (IM) as was used clinically in the trials for Glybera^12^; g) a low dose of 5e10 vg/mouse was chosen as this is sufficient to provide adequate FLuc signal but not so high that the maximal expression would already be reached thereby masking potential potency differences. Twelve mice (6 male, 6 female) were injected IM and live imaged weekly for FLuc expression (**Figures 4e, S12a**). Time course results revealed that human-rAAV1 achieved significantly higher skeletal muscle expression than baculo-rAAV1 in both sexes (*P <* 0.0001 at week 8) (**Figures 4g, S12c,d**).

To determine whether potency differences were restricted to IM delivery of rAAV1, we repeated the experiment with rAAV8—a serotype currently being tested as an intramuscular HIV passive vaccine vector^34^, despite reports of poor transduction in human skeletal muscle^35^ and other tissues^36,37^, as well as high neutralizing antibody levels in HIV+ individuals^38^. Here again, 12 mice (6 male, 6 female) were injected IM with 5e10 vg of ssAAV8-CMV-FLuc vector and live imaged weekly for FLuc expression (**Figures 4f, S12b**). Human-rAAV8 achieved significantly higher expression than baculo-rAAV8 in both sexes (*P* <0.0001 at week 8) (**Figures 4h, S12c,d**).

### Significant differential potency and sexually dimorphic functional transduction was observed with rAAV in mouse liver in vivo

Administration of rAAV IM is typically only done when treating muscle disorders or when using muscle as a secretion depot for passive vaccines^35^. Many gene therapy trials administer vector intravenously (IV) to facilitate distribution to internal organs not easily accessible by direct injection. Thus, we next assessed whether the potency differences seen with IM administration would be recapitulated with IV administration of human and baculovirus-*Sf9* produced rAAV. For this deeper set of comparisons, we utilized vector lots produced from a single facility and our custom dual-use transfer vector (**Figure S1**) to even more stringently control for variations between the two production platforms. Twenty-four mice (12 male, 12 female) were injected IV with 5e10 vg/mouse of ssAAV1-EF1α-FLuc produced in either the human or baculovirus-*Sf9* platform and live imaged weekly for FLuc expression (**Figure 5a**). Results highlighted several key findings. First, human-rAAV1 achieved significantly higher FLuc expression than baculo-rAAV1 (*P* < 0.005-0.0001 at week 4) (**Figure 5c**). Second, significant sexually dimorphic differences were seen where males experienced greater functional liver transduction than females (*P* < 0.04-0.0001). While sexually dimorphic liver transduction has been observed previously with human-produced rAAV^39^, we believe this is the first observation of this phenomenon with baculovirus-*Sf9* rAAV.

**Figure 5.**
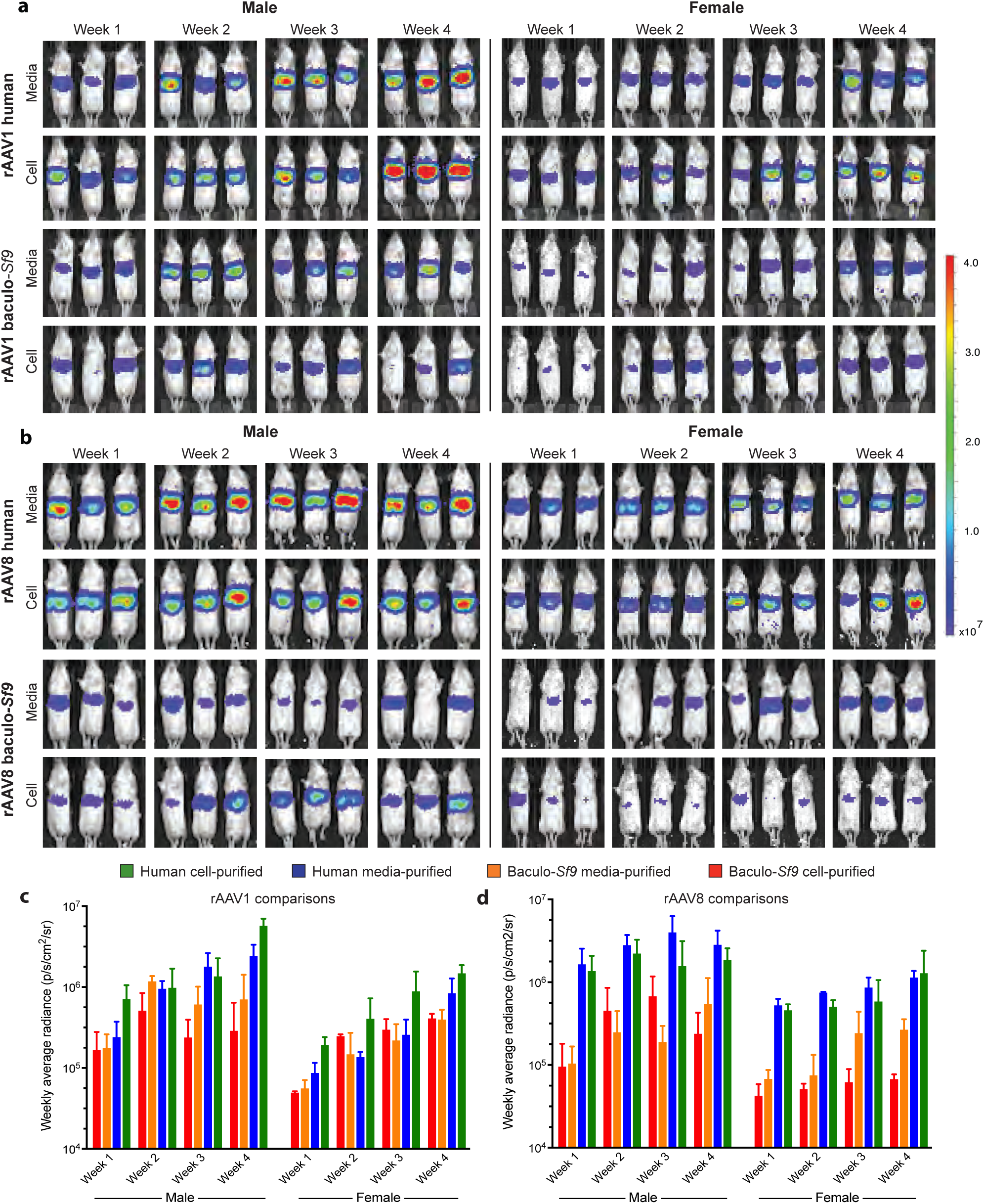
Human-produced rAAV produced significantly more functional liver transduction *in vivo* following intravenous administration and functional transduction is sexually dimorphic with rAAV from both platforms. **(a)** *In vivo* time course functional transduction assays comparing human and baculovirus-*Sf9* produced ssAAV1-EF1α-FLuc after IV tail vein administration (5E10 vg/mouse) in age-matched siblings. Mean radiance (p/s/cm^2^/sr) displayed with all mice imaged on their ventral side on the same shared scale. **(b)** Same as (a) but with rAAV8; same shared scale as (a). **(c)** Quantification of (a). Human rAAV1 is more potent than baculovirus-*Sf9* rAAV1 (*P* < 0.005-0.0001 at week 4) and males have higher functional liver transduction than females (*P* < 0.04-0.0001). **(d)** Quantification of (b). Human rAAV8 is also more potent than baculovirus-*Sf9* rAAV8 (*P* < 0.03-0.0001) and males again have higher functional liver transduction than females (*P* < 0.009-0.0001). Detailed statistics in Table S23.

To determine whether the significant functional and sexually dimorphic hepatic expression differences were restricted to rAAV1, we repeated the same *in vivo* experiment with rAAV8. Here again, 24 mice (12 male, 12 female) were injected IV with 5e10 vg/mouse of ssAAV8-EF1α-FLuc produced in either platform and live-imaged weekly for FLuc expression (**Figure 5b**). The results demonstrated that our previous differential potency findings with IV rAAV1 were not serotype specific, as they replicated with rAAV8 (*P* < 0.03-0.0001) (**Figure 5d**). Significant sexual dimorphism was again observed, with males having significantly greater functional liver transduction than females (*P* < 0.009-0.0001).

### Human-rAAV has significantly greater potency in human liver in vivo

To determine whether the significant potency differences seen in mouse liver following IV administration would also be seen in humans, we repeated the same *in vivo* experiment but now using humanized liver mice. Sixteen humanized liver mice (8 male, 8 female) were produced where male mice were transplanted with human male hepatocytes and female mice were transplanted with human female hepatocytes to >80% human liver repopulation (**Figure 6a, Table S24**). All humanized liver mice were injected IV with 5e10 vg/mouse of ssAAV1-EF1α-FLuc produced in either platform and live imaged for FLuc expression (**Figure 6b**). Results demonstrated that human-produced rAAV1 achieved significantly higher functional human liver transduction than baculovirus-*Sf9* produced rAAV1 (*P* < 0.005 at week 8) in male humanized mice. The same trend was observed in the female cohort, albeit with a higher P value (*P <* 0.08 at week 8) due to decreased power from the loss of two female subjects during the experiment (one female died post-transplant pre-AAV administration and one female had a failed tail vein injection of AAV). These results suggest that our previous findings with IV administered rAAV in mouse liver were not species-specific, as they were replicated in human liver *in vivo* as well. Note, these experiments were not repeated with rAAV8 given previous studies demonstrating that rAAV8 does not functionally transduce human liver *in vivo*^36,37,40-42^.

**Figure 6.**
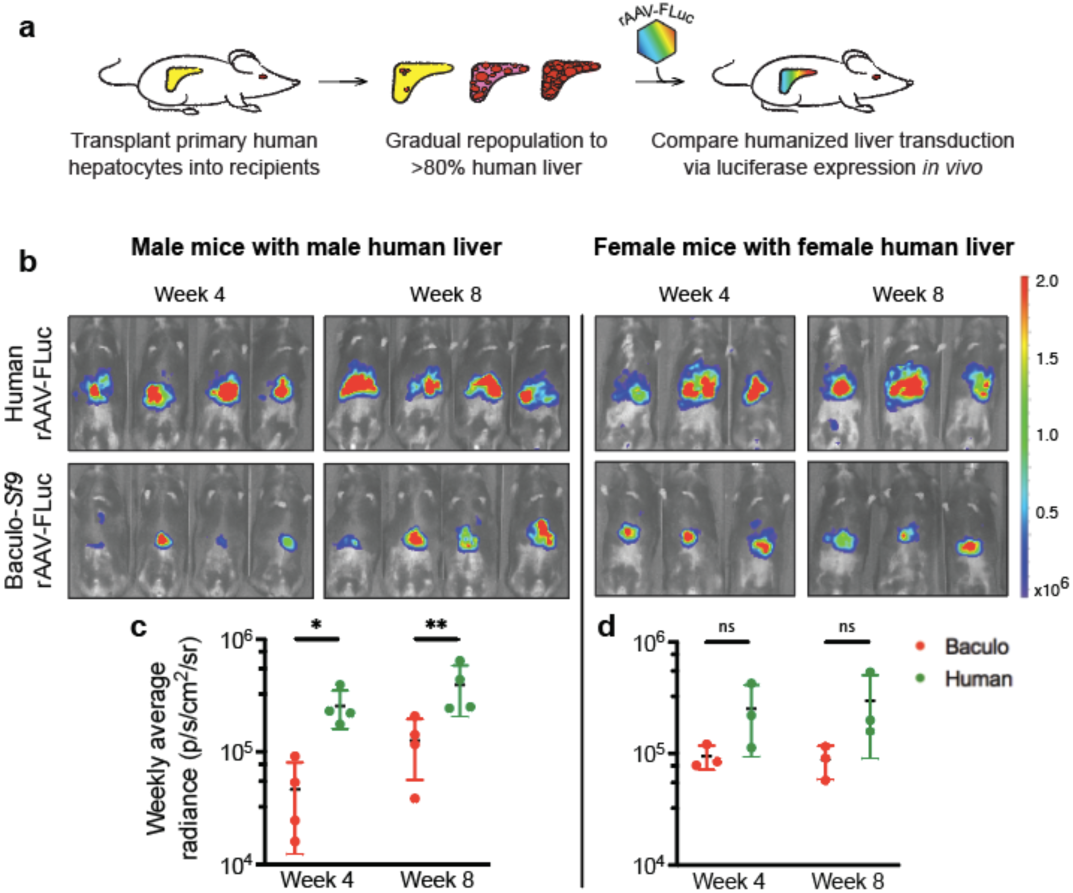
rAAV produced using the human manufacturing platform has significantly better functional human liver transduction *in vivo*. **(a)** Schematic illustrating the production of humanized liver mice used for assessing comparative functional human liver transduction *in vivo* with rAAV expressing Firefly Luciferase (FLuc). **(b)** *In vivo* time course functional transduction assays comparing human and baculo-*Sf9* produced ssAAV1-EF1α-FLuc after IV tail vein administration (5E10 vg/mouse) in age-matched humanized liver mice. Mean radiance (p/s/cm^2^/sr) displayed with all mice imaged on their ventral side on the same shared scale. **(c)** Quantification of rAAV1 FLuc radiance in male human liver. Each symbol = mean signal (+/- SD) from 4 humanized mice. **(d)** Quantification of rAAV1 FLuc radiance in female human liver. Each symbol = mean signal (+/- SD) from 3 humanized mice. * *P* ≤ 0.05, ** *P* ≤ 0.01, *** *P* ≤ 0.001, **** *P* ≤ 0.0001.

## Discussion

Of the nearly 400 FDA-approved protein-containing therapies in use in the United States—enzymes, growth factors, interferons, hormones, blood products, vaccines, antibodies, anti-venoms, anti-toxins, immune globulins, gene therapies, cell therapies, etc.—only two (both vaccines) are produced in any insect manufacturing system (**Table S25**)^43,44^. Within the rAAV gene therapy space, the human and insect manufacturing systems are predominantly used for production. Two prior studies comparing vectors produced in human and insect cells have been published by groups who were involved in developing the insect system^45,46^, and we believe that our study is the first independent comparative analysis of vectors from the two systems.

Clinicians, patients and regulators need to be confident in the safety and efficacy of rAAV for gene delivery. One safety concern is the potential for immunotoxicity against insect impurities^47^. Indeed, several recent severe adverse events with fever^8,9^ occurred following high dose rAAV administration with baculovirus-*Sf9* produced vector. Given the recent explosion of rAAV use--in trials for gene therapy, passive vaccines, and as a critical delivery agent in the rapidly expanding gene editing space--a thorough characterization of rAAV produced using different manufacturing methods was warranted.

Here we summarize our discoveries and those findings with critical implications for the field of gene therapy: when given at the same vector genome dose, human-produced rAAVs are more potent than baculovirus-*Sf9* rAAV vectors both *in vitro* and in numerous mouse and human tissues *in vivo*; rAAV capsids can be post-translationally modified (acetylation, methylation, phosphorylation, O-glcNAcylation) and these capsid PTMs differ when produced in the baculovirus-*Sf9* and human production platforms; HCP impurities differ in vector lots between platforms and can have their own PTMs, including potentially immunogenic N-linked glycans; capsid PTMs and HCP impurities were seen across all rAAV serotypes, manufacturers, purification types and different generations of the manufacturing technologies; packaged rAAV genomes are epigenetically methylated in both production platforms and baculovirus-*Sf9* rAAV has significantly more methylation in key repressive regions; lastly, regardless of the manufacturing platform, functional rAAV transduction is sexually dimorphic in the liver when administered intravenously.

It is known that human and insect cells have different capacities to post-translationally modify and fold proteins^10^. However, we were initially hesitant to speculate on rAAV capsid PTM presence given a publication from 2006 stating that there were none^48^. We noted several key differences between this study and ours. First, they used wild-type AAV2 produced in HeLa cells with wild-type replication-competent Adenovirus-2. Modern rAAV production now produces vector in HEK293 cells and provides the adenoviral helper function from Adenovirus-5 *in trans* from plasmids^4^. Collectively, the differences in cell type (HeLa vs. HEK293), method of helper viral administration (live Ad-2 vs. Ad-5 plasmid), and type of AAV (wtAAV vs rAAV), made us question whether the AAV particles each of us assessed were comparable. Indeed, with these differences and using more sensitive LC-MS/MS instruments, we observed numerous rAAV capsid PTMs. Additionally, there have been several recent attempts^16,45,46,49^ to characterize rAAV vector lots, however each had limitations in technology or scope, and called for the deeper research performed here. Our study used orthogonal methods to deeply characterize and validate differences in a sensitive and unbiased manner.

Human HEK293 and insect *Sf9* cells are both capable of methylating recombinant DNAs produced within them, given that each expresses the known methyltransferase machinery DNMT1 and DNMT3. However, no gene therapy literature existed on whether rAAV genomes are methylated when produced in either cell line. In eukaryotes, cytosine methylation is functionally used to regulate genomic expression. Promoter methylation, like we observed on baculovirus-*Sf9* produced rAAV, is typically used to transcriptionally repress the associated gene^50, 51^. In contrast, intragenic methylation, like we observed on human-produced rAAV, is typically associated with active transcription^50, 52, 53^. Thus, our data both support that rAAV genomes can be methylated during production, and that they are differentially methylated in key regions depending on the platform used to produce them with human-produced rAAVs having a more favorable methylation pattern for vector potency.

As we designed our experiments, we had to consider limitations as to which experimental parameters we could control. First, while clinical trials use Good Manufacturing Practices (GMP) grade vector, it is not possible to acquire aliquots of existing GMP lots as an external investigator, and the cost to produce even a single GMP lot renders cross comparative paired studies as performed here, prohibitive if not impossible. Thus, although we used preclinical grade vector, we ensured that it was of the highest possible purity and was identically purified for each production platform. However, it is important to recognize that GMP vectors may or may not exhibit the differences we observed here.

Several technical challenges prevented us from performing desired enzymatic experiments to add/remove PTMs to intact capsids to assess their contributions to potency. For example, there are no pan-demethylase or pan-deacetylase enzymes that work on intact proteins. Similarly, deglycosidases like PNGase-F perform optimally on denatured proteins, making it difficult to assess whether capsid glycans influence vector potency without altering capsid integrity. Further, many of these modifications are on the lumenal face, and are not accessible. One could set up a number of mutational analyses knocking out each host gene encoding each respective transferase for each PTM; however, given the number of different PTMs, there are too many genes to knock out simultaneously without negatively affecting host cell viability.

Although we observed differences in PTMs on baculovirus-*Sf9* and human-produced vector lots by LC-MS/MS, it is important to note that PTM absence does not indicate that one was not present. Additionally, observed PTMs were substoichiometric, with the majority of identified peptides being unmodified. Further, labile modifications may be lost during sample preparation or ionization prior to detection. Quantifying PTM frequency is challenging. In many cases, PTMs are not site-localizable, which can dramatically sway accuracy of quantitative evaluations of site occupancy. Additionally, different peptides from an individual protein can differ in cleavage efficiency. Thus, the list we have reported here should not be considered exhaustive, but rather, partially complete. A requirement for inclusion in our list of observed PTMs was that the modification must be site-localizable. To ensure our spectra were correctly assigned via automated database searching, every spectrum matching to a post-translationally modified peptide was also manually validated.

The field would benefit from a continued examination of the influence of production variables on vector lot composition, stability, potency and safety. These include factors such as suspension versus adherent cell culture, different purification methods, vector age, and more. For example, what effect does the use of live viruses like baculovirus (or herpes simplex virus) have during production? The initial motivation to adopt the baculovirus-*Sf9* production system stemmed from poor vector yields in adherent HEK293 cells. However, yields from new suspension-adapted HEK293 cells boost vector yields up to 1E5 vector genomes (vg)/cell^24^, higher than that achieved with the baculovirus-*Sf9* system at 7E4 vg/cell^5–7^. Additional parameters that could be enhanced include improving continuous rAAV collection from suspension media^54^, boosting host expression of key production factors^55^, engineering other mammalian lines to safely and efficaciously produce at high yield, developing tunable cell-free methods to produce rAAV, and beyond. Given the catalog of observed differences between the two rAAV vector production platforms characterized herein, a continued investigation into the implications of these differences for the clinical and research communities is warranted.

## Supporting information

Supplemental Figures and Small Tables

**This article contains supporting figures and tables online.**

## Author Contributions

N.K.P. conceived the study. N.K.P, N.G.R., S.A.M., and A.P.M. designed experiments. N.K.P, N.G.R., S.A.M., N.P., L.H.M., G.C., C.A., R.D.L., D.T., S.S., K.H., P.S. and A.P.M. generated reagents, protocols, performed experiments, and analyzed data. N.K.P. wrote the manuscript. N.G.R., S.A.M., N.P., L.H.M., and N.K.P. generated the figures. All authors reviewed, edited and approved the manuscript.

## Acknowledgments

The authors wish to acknowledge the generous gift of 16 humanized liver FRG mice from Yecuris Corporation (www.yecuris.com); Lutz Froenicke and Emily Kumimoto of the UC Davis Genome Center for help with methylation sequencing; Jie Lie and Matthew Settles of the UC Davis DNA Technologies Core for help with WGBS data analysis; John Perrino of the Stanford University Cell Sciences Imaging Facility and Tom Moninger of the University of Iowa Central Microscopy Research Facility for help with TEM images; Yael Rosenberg-Hasson of the Stanford Human Immune Monitoring Core for help with the Luminex assays; Parastoo Azadi of the Complex Carbohydrate Research Center at the University of Georgia for glycoproteomic help; the University of Massachusetts for the baculovirus-*Sf9* AAV2-hAADC samples; Linda Couto of The Children’s Hospital of Philadelphia for Huh7 cells; Donghui Wang of UCSF for mouse tail vein injections; the University of Iowa Viral Vector Core Facility for collaborative viral productions; Bruce Conklin of the Gladstone Institute and UCSF for the human iPSC cells; the authors also wish to thank the following investigators for initial discussions: Carolyn Bertozzi and Mark Kay of Stanford University and Joseph DeRisi of UCSF. This research was supported by grants to N.K.P. from the NIH (K01-DK107607, U01-HL145795), an American Society of Gene & Cell Therapy Career Development Grant, the Sandler Family Foundation, a Stanford University Immunity, Transplantation and Infection Young Investigator Award, and a Vincent Coates Foundation Mass Spectrometry Laboratory seed grant; grants to the University of Iowa Viral Vector Core from the NIH (P01-HL51670), the Center for Gene Therapy of Cystic Fibrosis (P30-DK54759), and the Holden Comprehensive Cancer Center (P30-CA086862); computing resources from Wynton a UCSF Shared Research Computing Cluster (https://wynton.ucsf.edu/); methylation sequencing was carried out at the DNA Technologies and Expression Analysis Core at the UC Davis Genome Center and supported by NIH Shared Instrumentation Grant 1S10OD010786-01; and funding from the Chan Zuckerberg Biohub and Cold Spring Harbor Labs; N.G.R. and S.A.M. were supported by the NIH (U01-CA207702) and HHMI, and S.A.M. was also supported by an NIH fellowship F32-GM126663; and N.P. was supported by an NIH Microbial Pathogenesis and Host Defense training grant T32-AI060537. The contents of this publication are solely the responsibility of the authors and do not necessarily represent the official views of the funding bodies, or respective universities, institutes and organizations.

## Materials and Methods

### Generation of pFB.AAV-EF1ɑ-FLuc-WPRE-hGHpA_BAC-293 dual-use transfer vector plasmid

pAAV-EF1ɑ-FLuc-WPRE-hGHpA transfer vector plasmid^37^ (Addgene Cat#87951) was cut with *SbfI* dual-cutter just beyond each ITR to create a 4.3-kb fragment. To prepare the universal shuttle backbone, pFB.AAV-msc_bGHpA #G0202 from the University of Iowa was also cut with *SbfI* and the resulting purified backbone was ligated to the 4.3-kb transfer vector fragment to create the 8.1-kb final construct. All components for insect replication (gentamicin-R and Tn7 transposition sites) and bacterial replication (ampicillin-R) are present on the dual-use backbone. Academic MTAs to Dr. Paulk at UCSF were acquired from The Salk Institute for WPRE use, and University of Iowa for pFB.AAV-msc_bGHpA plasmid backbone. The final pFB.AAV-EF1ɑ-FLuc-WPRE-hGHpA_BAC-293 plasmid has been deposited at Addgene (Cat#118412). Figure S1a was made with SnapGene v4.2.11; S1b was made with GraphPad Prism v8.0.1.

### Plasmid transfection and Firefly Luciferase expression assessment to validate pFB.AAV-EF1ɑ-FLuc-WPRE-hGHpA_BAC-293

Each well of a 6-well plate was seeded with 500K HEK293T cells (ATCC Cat#CRL-3216). Once cells reached 80% confluency, equimolar concentrations of plasmid (custom plasmid and control plasmids) were transfected via Ca_3_(PO_4_)_2_ transient transfection and media was changed after 8-hr. Cells were lysed 48-hrs later for FLuc quantitation using a Luciferase 1000 Assay System kit (Promega Cat#E4550) according to manufacturer’s instructions and read on a Veritas luminometer. Experiments were performed in technical triplicate.

### Generation of rAAV bacmids for baculovirus production

A custom pFB.AAV-EF1ɑ-FLuc-WPRE-hGHpA bacmid was produced using the Bac-to-Bac production system where MAX Efficiency DH10Bac *E. coli* (Thermo Cat#10361012) contained a baculovirus shuttle vector that can recombine with a donor plasmid, pFastBac (pFB), to create an expression bacmid containing the cloned construct for eventual rAAV transfer vector production. The donor plasmid was the custom pFB.AAV-EF1ɑ-FLuc-WPRE-hGHpA_BAC-293 construct described above. 10-ng of pFB.AAV-EF1ɑ-FLuc-WPRE-hGHpA_BAC-293 donor plasmid was transposed into 100-mL DH10Bac competent cells and plated on DHBac10 LB agar plates for 48-72 hours at 37°C in a non-CO_2_ incubator. Plates with colonies were kept at 4°C thereafter until ready for use. A single large isolated transposed white colony was picked and re-streaked for isolation on another DHBac10 LB agar plate. The confirmation colony was inoculated into DHBac10 LB broth and the bacmid grown for 20-hrs in a 200-rpm shaking incubator at 37°C. Bacmid DNA was isolated with a NucleoBond Xtra Midi EF kit (Macherey-Nagel Cat# 740.420.50), resuspended in TE, and stored in aliquots at -20°C. The packaging bacmids for generating baculoviruses for producing different serotypes of rAAV were produced as described above, except using pFB donor plasmids encoding AAV2 Rep and the capsid of interest (pFB-AAV-Rep2/Cap8 or pFB-AAV-Rep2/Cap1) from the University of Iowa.

### Generation of baculoviral lots for rAAV production

For BAC-AAV-EF1ɑ-FLuc-WPRE-hGHpA and BAC-AAV-Rep2/Cap1 or 8, 2×10^6^ *Sf9* cells (Expression Systems Cat# 94-001S) were seeded per well in a 6-well plate in protein-free ESF 921 media (Expression Systems Cat# 96-001) with no additives or antibiotics, using cells from a 3-day-old suspension culture in mid-log phase (95-100% visual viability). Cells were allowed to attach at 27°C for 1-hr (∼80% confluency). The transfection solution was prepared with 45-mg of purified bacmid in 600-mL media, and mixed with 36-mL of Cellfectin (Invitrogen Cat#10362-010) in 600-mL media, mixed and incubated at RT for 30-min. Adherent cells were washed and then transfected with 700-mL per well of transfection solution diluted in 3-mL media. Cells were incubated for 5-hr in a 27°C incubator in a humidity chamber. Following incubation, transfection mix was removed and 2-mL of fresh media was added per well before incubation for 5 days at 27°C. Resultant baculovirus was harvested from the media supernatant and mixed with serum to [final] 5% for storage. Media was spun at 500xG for 5-min to pellet any cellular debris. Solution was filtered through a Millex 0.22-µm sterile syringe filter with PES membrane (EMD Millipore Cat# SLGP033RB) and aliquoted. Baculoviral titers were determined using a BackPak Baculovirus Rapid Titer kit (Clontech Cat# 631406).

### Production of rAAV from baculovirus infection of *Sf9* cells for paired productions

For rAAV production, 2×10^6^ suspension *Sf9* cells/mL were seeded in 400-mL ESF 921 media in roller bottles. 24-hrs later, cells were counted for infection calculations and to ensure high viability. 100-mL of fresh media was added to each flask prior to infection. Baculoviral aliquots were thawed, resuspended, and 0.5-MOI of each baculovirus (transfer vector virus and pseudotyping virus) were added directly to the media and bottles were returned to the incubator for 72-hr at 27°C. For production of the special empty capsid lots, the transfer vector baculovirus was omitted and only the pseudotyping baculovirus was used to ensure no transfer vector was present to package.

### Production of rAAV in HEK293FT cells for paired productions

Recombinant AAV vector productions were produced in 40 15-cm plates using a Ca_3_(PO_4_)_2_ transient triple transfection protocol in adherent HEK293FT cells (Thermo Fisher Cat# R70007) in DMEM + 10% FBS +1% penicillin/streptomycin. Plasmids included: pHelper (1000-mg/40 plates), pFB.AAV-EF1ɑ-FLuc-WPRE-hGHpA_BAC-293 transfer vector plasmid (500-mg/40 plates), and a pseudotyping plasmid (pRep2/Cap8 or pRep2/Cap1; 1500-mg/40 plates). Transfection solution was incubated with cells for 16-hr at 37°C, then the media was changed to serum-free Expi293 media (Thermo Cat#A14351) for the remaining 60-hr incubation at 37°C in 5% CO_2_ for a total incubation of 76-hr. For production of the special empty capsid lots, the transfer vector plasmid was omitted and only the pHelper and pseudotyping plasmids were used to ensure no transfer vector was present to package.

### Universal harvest protocol for isolating rAAV from cell lysates and media supernatant from both human and baculovirus-*Sf9* production platforms for paired productions

To isolate rAAV particles from cell lysates and media supernatant, the following protocols were followed. For cell-purified rAAV: cell pellets were lysed with a 12-min glass bead vortex, treated with 100-U/mL Benzonase (Sigma Cat# E8263-25KU) for 30-min at 37°C followed by addition of 10% sodium deoxycholate and another 30-min 37°C incubation to dissociate particles from membranes. Then they were iodixanol gradient-purified in OptiSeal polyallomer tubes (Beckman Cat# 362183) in a VTi-50 rotor spun at 40,000-rpm for 1-hr 50-min at 18°C in an Optima LE-80K-Beckman ultracentrifuge, further purified by MustangQ ion exchange chromatography on a peristaltic pump, and buffer exchange through an Amicon Ultra-15 centrifugal filter concentrator (Millipore Cat#UFC910096) into 1X dPBS pH 7.4 + 180-mM NaCl + 0.001% Lutrol (v/v). For media-purified rAAV: media was first put through a 100-kDa MWCO filter (Spectrum Cat#S02-E100-10N), then tangential flow filtered at 400mL/min with TMP of 4.0, then Triton-X-100 was added to 0.5% [final] and incubated in a shaking water bath at 37°C for 1-hr, solution was then treated with 200U/l Turbonuclease (Sigma Cat#T4330-50KU) for 1-hr at 37°C, then iodixanol gradient purified as described above, followed by MustangQ ion exchange chromatography as described above, and buffer exchanged as described above. rAAV vectors were simultaneously titered by TaqMan qPCR (on the same plate) within the hGHpA element with the following primer/probe set:

Fwd: 5’-TGTCTGACTAGGTGTCCTTCTA-3’

Rev: 5’-CTCCAGCTTGGTTCCCAATA-3’

Probe: 5’/6-FAM/AAGTTGGGAAGACAACCTGTAGGGC/IBFQ/-3’

### Production of rAAV in HEK293T cells for non-paired productions

Recombinant AAV vector productions were produced using a Ca_3_(PO_4_)_2_ transient triple transfection protocol in adherent HEK293T cells (ATCC Cat#CRL-3216) followed by double cesium chloride density gradient purification and dialysis as previously described^35,38^, and resuspended in dPBS with 5% sorbitol (w/v) and 0.001% Pluronic F-68 (v/v). Plasmids included: pAd5 helper, rAAV transfer vector (ssAAV-CAG-TdTomato, Addgene #59462), and pseudotyping plasmids for each capsid of interest. rAAV vectors were titered by TaqMan qPCR within the CAG promoter with the following primer/probe set:

Fwd: 5’-GTTACTCCCACAGGTGAGC-3’

Rev: 5’-AGAAACAAGCCGTCATTAAACC-3’

Probe: 5’/FAM/CTTCTCCTCCGGGCTGTAATTAGC/BHQ-1/-3’

### Two-dimensional gel electrophoresis

Protein samples for each production platform were fractionated by two-dimensional gel electrophoresis. To determine whether differences between gels resulted from differences in capsid PTMs, half of the samples were treated with a broad-spectrum lambda protein phosphatase (NEB Cat#P0753L) and PNGase-F (Promega Cat#V4831) prior to two-dimensional gel electrophoresis. 50-mg total protein from each production platform was used per preparation. Total sample volumes were adjusted to 200-μL after addition of 20-μL each of the 10X MgCl_2_ and 10X phosphatase buffer (supplied with enzyme) per tube (regardless of enzyme treatment). 1.25-μL of enzyme was added to the phosphatase-treated samples, with the remainder being water. Samples were gently mixed and incubated for 2-hr at 30°C at 1,000-rpm. 100-mM Dithiothreitol (DTT) in water (BIO-WORLD Cat#404200001) was then added to each tube to a final concentration of 5-mM, and samples were heated at 95°C for 5-min. Samples were cooled to 25°C before adding 5-μL of PNGase-F and incubated at 37°C overnight at 1,000 rpm. To separate proteins by their isoelectric points, samples were then loaded in equal volumes into 24 wells of an Agilent 3100 OFFGEL Fractionator using a high-resolution fractionation kit (Agilent Cat#5067-0201). Each sample, with and without enzyme treatment, was run on a separate lane of a 24 well gel strip with a linear gradient from pH 3-10 following manufacturer’s instructions. Runs were completed upon reaching a total of 64-kW hrs. Samples from each well were concentrated to ∼30-μL with Amicon Ultra 10-KDa molecular weight cutoff spin filters (EMD Millipore Cat#UFC501008) for 2-hrs before loading into a 4-12% Bis-Tris SDS-PAGE gel (Invitrogen Cat#NP0322BOX). Gels were washed, fixed, and stained overnight with SYPRO Ruby protein gel stain (FisherSci Cat#S12000) following the manufacturer’s instructions. Gels were then washed and imaged with a UV transilluminator.

To test whether proteases were present (in either the rAAV lots, phosphatase or PNGase-F reagents) and responsible for the banding patterns seen in Fig. S6, samples were prepared as above, but with or without pre-treatment with protease inhibitor (Roche Cat#04693159001) in combination with phosphatase treatment, PNGase-F treatment, or both. Lactoferrin was used as a positive glycoprotein control.

### Subsequent Western blots on 2D-gel electrophoresis SDS-PAGE gels

Gels were transferred onto nitrocellulose membranes (Bio-Rad Cat#1620112) with a Trans-Blot Turbo Transfer System (Bio-Rad), blocked with Odyssey Blocking Buffer (TBS) (LI-COR Cat#927-50000), blotted with rabbit anti-AAV-VP1/2/3 polyclonal antibody (ARP Cat#03-61084) at 1:200 overnight at 4°C, then washed with PBS-T and incubated with secondary goat anti-rabbit IgG IRDye 800CW (LI-COR Cat#926-32211) at 1:10,000 for 1-hr at 25°C. The membranes were then washed with PBS-T and visualized on a Licor Odyssey CLx Infrared Imaging System. Imaging analysis was performed using the Licor Image Studio Lite software.

### Proteolytic digests of rAAV samples for LC-MS/MS

Protein preparations for paired rAAV productions were done using equal amounts of total protein (10-50 μg depending on the run). Each sample was precipitated using 4X volume of LC-grade acetone at -80°C overnight. Precipitated protein was centrifuged at 12,500xG for 15-min at 4°C. Supernatant was decanted and discarded and the protein pellet dried within a biosafety hood for 30-min. The pellet was resuspended in 100-μL of 0.2% Protease Max Surfactant Trypsin Enhancer (Promega Cat# V2071), 50-mM ABC, and reduced using DTT to a final concentration of 10-mM at 55°C for 30-min. Following reduction, proteins were alkylated with 20-mM propionamide at 25°C for 30-min. For those samples undergoing Endoglycosidase-H (Endo-H) treatment, samples were heated to 95°C for 5-min, followed by the addition of 5-μL Endo-H reaction buffer and 5-μL Endo-H (Promega Cat#PRV4875, 2,500-U). Deglycosylation proceeded for 4-hours at 37°C. Endo-H is a recombinant glycosidase which hydrolyses the bond connecting the two GlcNAc groups comprised in the chitobiose core, leaving a single GlcNAc covalently bound for mass spectrometry detection. 500-ng of Trypsin/Lys-C Mix (Promega Cat#V5072) was added and allowed to digest overnight for ∼18-hr at 37°C. The digest was acidified to 1% formic acid to quench protease activity. Peptides were purified on C18 MonoSpin SPE columns (GL Sciences Cat#5010-21701), and peptide pools dried to completion in a speed vac.

### Liquid chromatography–mass spectrometry for non-glycosylated peptides

Peptide pools were reconstituted in 0.2% formic acid, 2% acetonitrile, and water and then injected onto a Waters M-Class UPLC. The analytical column was pulled and packed in-house using a C18 matrix (Dr. Maisch, 2.4-μM, Pur), at approximately 25-cm in length. The flow rate was 300-nL or 450-nL per minute where mobile phase A was 0.2% formic acid in water, mobile phase B was 0.2% formic acid in acetonitrile. The mass spectrometer used was an Orbitrap Fusion Tribrid (Thermo Scientific) set to acquire data in a dependent fashion using multiple fragmentation types: collision-induced dissociation (CID), higher-energy collisional dissociation (HCD), and electron-transfer dissociation (ETD). The instrument parameters used in this study were the following: CID data were generated when the precursor mass resolution was 120,000 with a *m*/*z* window of 400-1500, and charge states of 2+, 3+ and 4+ were sampled. The precursor automated gain control (AGC) settings were 200,000 ions and the “fastest” mode was used for MS/MS in the ion trap. The ion trap sampled the most intense precursors where the isolation window was set at 1.6-Da and the collision energy at 35. In a HCD experiment, the settings were the same as for CID except a normalized collision energy of 42 was used and fragment ions were sent to the Orbitrap for detection. Finally, in ETD/HCD triggered experiments, the precursor mass resolution was 120,000, with a *m*/*z* window of 400-1600, and charge states of 2-6+ were sequenced and precursor AGC target was 400,000. The instrument alternated sequencing the same precursor masses first by ETD--where the settings were a 50-ms reaction time and 200,000 reagent target--and a max ETD time of 200-ms. Fragment ions were read out in the ion trap in a centroid fashion. HCD had a collision energy of 32 and a maximum inject time of 54-ms where fragment ions were read out of the Orbitrap in a centroid fashion. The isolation window for both the fragmentation types was 1.2-Da and the data dependent acquisition exclusion settings allowed for sampling of the same precursor three times before it was placed on the exclusion list. The exclusion list was set to 10-ppm and in the ETD/HCD trigger experiment timed out after 10-sec.

### Triggered liquid chromatography–mass spectrometry for glycosylated peptides

Samples were analyzed by online nanoflow LC-MS/MS using an Orbitrap Fusion Tribrid mass spectrometer (Thermo Fisher Scientific) coupled to a Dionex Ultimate 3000 HPLC (Thermo Fisher Scientific). Sample were loaded via autosampler onto a 20-µL sample loop and injected at 0.3-µL/min onto a 75-µm x 150-mm EASYSpray column (Thermo Fisher Scientific) containing 2-µm C18 beads. Columns were held at 40°C using a column heater in the EASY-Spray ionization source (Thermo Fisher Scientific). Samples were eluted at 0.3-µL/min using a 90-min gradient and 185-min instrument method. Solvent A was comprised of 0.1% formic acid in water, whereas Solvent B was 0.1% formic acid in acetonitrile. The gradient profile was as follows (min:%B) 0:3, 3:3, 93:35, 103:42, 104:98, 109:98, 110:3, 185:3. The instrument method used an MS1 resolution of 60,000 at FWHM 400-*m/z*, an AGC target of 3e5, and a mass range from 300 to 1,500-*m/z*. Dynamic exclusion was enabled with a repeat count of 3, repeat duration of 10-sec, and an exclusion duration of 10-sec. Only charge states 2-6 were selected for fragmentation. MS2s were generated at top speed for 3-sec. HCD was performed on all selected precursor masses with the following parameters: isolation window of 2-*m/z*, 28-30% collision energy, orbitrap (resolution 30,000) detection, and an AGC target of 1e4 ions. ETD was performed if: (a) the precursor mass was between 300 and 1000-*m/z*; and (b) 3 of 7 glyco-fingerprint ions (126.055, 138.055, 144.07, 168.065, 186.076, 204.086, 274.092, 292.103) were present at +/- 0.5-*m/z* and greater than 5% relative intensity. ETD parameters were as follows: calibrated charge-dependent ETD times, 2e5 reagent target, and precursor AGC target 1e4.

### Mass spectra data analysis

MS/MS data were analyzed using both Preview and Byonic v2.10.5 software (ProteinMetrics). Data were first analyzed in Preview to validate calibration and trypsin/chymotrypsin digestion efficiency. Initial Byonic analyses used a concatenated FASTA file containing the AAV sequences, host proteomes (*Spodoptera frugiperda, Homo sapiens*, and *Autographa californica multiple nuclepolyhedrovirus* (baculovirus)) and other likely contaminants and impurities to verify sample purity. Once sample complexity was determined, a second round of Byonic analyses were completed using a targeted FASTA file containing the AAV sequence of interest and likely impurities to identify peptides and potential PTMs. Data were searched at 10-12 ppm mass tolerances for precursors (depending on the run), with 10-12 ppm or 0.1-0.4 Da fragment mass tolerances for HCD and ETD fragmentation, respectively. The algorithm allowed up to two missed cleavages per peptide and semi-specific, N-ragged tryptic digestion. These data were validated at a 1% false discovery rate using standard reverse-decoy techniques^56^. The resulting identified peptide spectral matches and assigned proteins were then exported for further analysis and validated using custom tools to provide visualization and statistical characterization. All PTM mass spectra were also manually validated.

### *De novo* glycan identification

Glycopeptide sequences were determined by *de novo* manual interpretation of HCD and ETD mass spectra. Extracted chromatograms were generated for all MS2s containing 204.0867 ions (+/- 10-ppm), and all spectra containing this ion were analyzed. HCD spectra were used to approximate glycan structures, and ETD spectra were used to define peptide sequence and site-localize modifications. GalNAc and GlcNAc were distinguished by their defined HexNAc fingerprints, as described previously^57^.

### *In vitro* transduction analysis by Firefly luciferase quantitation

Five cell types were assessed: immortalized human HEK293T cells (ATCC Cat# CRL-3216), immortalized mouse C2C12 myoblasts (ATCC Cat#CRL-1772), immortalized human Huh7 hepatocarcinoma cells (gift of Linda Couto), primary Hs27 human fetal foreskin fibroblasts (ATCC Cat#CRL-1634) all maintained in DMEM (Gibco Cat# 11995065) supplemented with 10% FBS and 1% antibiotic/antimycotic; and primary human iPSC cells (Coriell Cat# GM25256) derived from the skin biopsy of a healthy 30-year-old Japanese male which were maintained with the following modifications from a recent protocol^58^: used Matrigel growth factor reduced basement membrane matrix (Corning Cat#354230) in feeder-free media conditions (Gibco Essential 8 Flex Media Cat# A2858501) with a seeding density of 35K/cm^2^. Accutase (STEMCELL Technologies Cat# 07920) and 10-M RHO/ROCK pathway inhibitor Y-276932 (STEMCELL Technologies Cat# 72304) were used to dissociate iPSCs to single cells during passaging. In all cell types, 1-hr prior to transduction at 80% confluency, media was changed, then 80K cells/condition were transduced with ssAAV1-EF1α-FLuc vectors diluted in dPBS at MOI 80K. The following day, media was replaced to remove vectors that did not transduce and media was not changed again. FLuc levels were measured 3-days post-AAV administration using a Luciferase 1000 Assay System (Promega Cat#E4550) per manufacturer’s instructions and read on a Veritas luminometer at: 100-μL luciferin injection, 2-sec delay, 10-sec measure. Experiments were performed in technical triplicate and corrected for background by subtracting signal from PBS-diluent negative control wells. Figure 4a was generated with GraphPad Prism v8.0.1.

### Human tissue

We obtained anonymized human induced pluripotent stem cells derived from normal controls enrolled in ongoing, CHR-approved studies for basic research purposes by the human gamete, embryo and stem cell research committee to Dr. Bruce Conklin under the UCSF IRB#10-02521 “Induced Pluripotent Stem Cells for Genetic Research”. The iPS cells were originally derived from a 30-year-old Japanese male who signed a consent form approving the donation of iPS cells to the public stem cell bank Coriell (Cat# GM25256).

### Mixed titration/empty spike-in transduction analyses *in vitro*

Human HEK293T cells were maintained as described above. One hour prior to transduction at 80% confluency, media was changed, and 60K cells were transduced with varying ratios of full:empty ssAAV1-EF1α-FLuc-WPRE-hGHpA vectors produced by either platform and diluted in dPBS (only cell-purified vector was used). The total capsid content was kept constant at MOI 30K, while the ratio of full:empty varied: 30K:0K (0% empty), 27K:3K (10% empty), 15K:15K (50% empty), 3K:27K (90% empty), 0K:30K (100% empty). The following day, media was replaced to remove non-transducing vectors and the media was not changed again. FLuc levels were measured 3-days post-AAV administration using a Luciferase 1000 Assay System kit (Promega Cat#E4550) per manufacturer’s instructions and read on a Veritas luminometer with settings: 100-μL luciferin injection, 2-sec delay, 10-sec measure. Experiments were performed in 6 biological replicates and corrected for background by subtracting the signal from the PBS-diluent negative control wells. For the spike-in experiment, experimental conditions were the same except the baculoviral full MOI was kept constant at 30K and additional empty vector was spiked-in from either platform at 10% (3K) or 100% (30K) and assessed as previously described. This experiment was a single biological replicate performed in technical triplicate and corrected for background by subtracting the signal from PBS-diluent negative control wells. Figures 4b-d were generated with GraphPad Prism v8.0.1.

### Live *in vivo* transduction analysis by Firefly luciferase imaging and quantitation in mice

All mice having received either intravenous normodynamic lateral tail vein injections or intramuscular (*tibialis anterior*) injections of 5E10 vg/mouse of ssAAV-EF1α-FLuc (both rAAV8 and rAAV1) were imaged non-invasively every 7-days on a Xenogen IVIS Spectrum imaging system (Caliper Life Sciences). D-luciferin substrate (Biosynth Cat#L-8220) was administered at 120-mg/kg in saline by intraperitoneal injection with a 1-cc insulin syringe. Images were acquired 10-min after luciferin administration under inhalation isoflurane anesthesia. Living Image v4.5 software was used for image analysis and average radiance was quantified in p/s/cm^2^/sr. All mice in each experiment are shown on the same non-individualized radiance scale to enable accurate comparisons of bioluminescent intensity.

### Silver staining

Viral lots were TaqMan qPCR titered (described above) and 1E10 vg worth of vector was mixed with 5-μL of 4X NuPage LDS Buffer (Invitrogen Cat#NP0007) and 2-μL of 10X NuPage Reducing Agent (Invitrogen Cat#NP0009) and boiled for 8-min. Boiled samples were loaded into 4-12% 1-mm 12-well Bis-Tris NuPage gels (Invitrogen Cat#NP0322PK2) along with 0.5-μL of Benchmark Unstained Protein Ladder (Invitrogen Cat#10747-012) and run in 1X NuPAGE MES buffer (Invitrogen Cat#NP0002) supplemented with 500-μL NuPage Antioxidant (Invitrogen Cat#NP0005) at 100-V for 2.8-hrs. Gels were rinsed in 18 MΩ water, fixed and stained using a SilverQuest Staining Kit (Invitrogen Cat#LC6070) according to the manufacturer’s instructions.

### Luminex cytokine assay

Primary human fetal foreskin fibroblasts (ATCC Cat#CRL-1634) were maintained in DMEM (Gibco Cat#11995065) supplemented with 10% FBS and 1% antibiotic/antimycotic and seeded at 12K per well in a 96 well plate. 24-hrs following seeding, at 24K confirmed cells/well (control wells for each condition were lifted, counted and verified for viability), media was changed and allowed to sit for 1-hr prior to transduction with four rAAV lots of ssAAV8-EF1α-Fluc-WPRE produced by either platform and purified from either cell lysate or media supernatant (as described above) at an MOI of 100K in a total final media volume of 200-µL. No further media changes or washes occurred until media harvest at 24-hrs following rAAV addition. Controls included ‘media only’ wells (no cells or rAAV), and ‘media + cells’ wells (no rAAV) for background normalization calculations. Media was harvested from each well, spun at 350-rcf for 5-min at 4°C to pellet any cells, and media supernatants were then aliquoted in four protein LoBind polypropylene tubes (Eppendorf Cat#022431081) per well for storage at -80°C until use in the assay. Conditions were set up in biological triplicate and run in technical duplicate. The assay was performed at the Human Immune Monitoring Center at Stanford University. A custom human 63-plex kit was purchased from eBiosciences and used according to the manufacturer’s recommendations with modifications described below. Briefly: beads were added to a 96-well plate and washed in a Biotek ELx405 washer. Thawed samples were added to the plate containing the mixed antibody-linked beads and incubated at 25°C for 1-hr followed by overnight incubation at 4°C with shaking. Cold and 25°C incubation steps were performed on an orbital shaker at 500-600 rpm. Following the overnight incubation, plates were washed in a Biotek ELx405 washer and then biotinylated detection antibody added for 75-min at 25°C with shaking. The plate was washed as above and streptavidin-PE was added. After incubation for 30-min at 25°C, a wash was performed as above and reading buffer was added to each well. Each sample was measured in duplicate. Plates were read using a Luminex 200 instrument with a lower bound of 50 beads per sample per cytokine. Custom assay control beads by Radix Biosolutions were added to all wells. Data were normalized to controls with no AAV added during production (plain host cells cultured for either 24-hr or 48-hr) and averaged across all biological replicates. Data were visualized in R Studio.

### Next-generation sequencing of packaged rAAV genomes using Fast-Seq

Total packaged gDNA was extracted from 1E11 full rAAV particles for each lot of rAAV8 and rAAV1 sequenced. AAV libraries were prepared following a novel Tn5 tagmentation-based protocol called Fast-Seq^33^. The following indexed adapters compatible with Illumina were obtained from IDT:

Index 1 (i7):

**CAAGCAGAAGACGGCATACGAGAT***NNNNNNNNNNNN*<u>GTCTC</u> <u>GTGGGCTCGG</u>

Index 2 (i5):

**AATGATACGGCGACCACCGAGATCTACAC***NNNNNNNNNNN N*<u>TCGTCGGCAGCGTC</u>

**bold** = P5/P7 adapter sequence

*italics* = unique 12-bp barcode index

<u>underline</u> = primer for mosaic ends added during tagmentation

Each adapter contained a 12-nucleotide unique barcode for identifying samples after multiplexing. Indexes used for each sample are listed in **Table S20**. The resulting library was diluted to 10-pM in 600-μL of HT1 hybridization buffer (Illumina Nextera XT kit Cat#FC-131-1024) and 10-μL was loaded onto a 300-cycle MiSeq Nano v2 flow cell (Illumina Cat#MS-102-2002) for paired-end 2 x 75-bp sequencing. Resultant reads were demultiplexed using Illumina’s bcl2fastq v2.19.0.316. Data were returned in fastq format and filtered using Trimmomatic^59^ to remove adapter sequences, low quality reads (PHRED score <30, or length <50-bp), unmapped and unpaired reads. Trimmed reads were then aligned to the rAAV transfer vector plasmid reference sequence with BWA v0.7.17^60^, using the mem algorithm. Alignments were saved as BAM files, which were then used to generate VCF files using GATK Haplotype Caller algorithm^61^. SNPs and indels identified in VCF files were filtered using BCFtools filter algorithm, with a 15X depth threshold and a 90% allele fraction requirement. A consensus sequence was generated using BCFtools consensus algorithm^62^. Alignment and fragment distribution statistics were obtained with Picard tools^63^. Coverage spanned 100% of the rAAV transfer vector genome reference sequence. Figure S11b was generated with ggplot. Figure S11c was generated with the ViennaRNA Web Services structure prediction software for DNA available at http://rna.tbi.univie.ac.at.

### Capsid protein thermal stability

Purified rAAV capsids of various serotypes produced by each manufacturing platform were compared for their denaturing thermal stability. 10-µL of undiluted rAAV of each lot was loaded into a high sensitivity capillary (Nanotemper Cat#PR-C006) and run on a Nanotemper Prometheus NT.48 nanoDSF instrument. Thermal denaturation programs were run with a 1°C/min thermal ramp from 30°C to 95°C with fluorescent detection at 330 and 350 nm. Data were analyzed with PR.ThermControl software v2.1.5 to determine the temperatures for T_onset_ and T_m_. Figures 1m, S7 and S8k were made with GraphPad Prism v8.0.1.

### Negative staining and TEM imaging

Grids were glow discharged in the presence of argon gas for 20-sec in a Denton Desktop Turbo III vacuum system and then covered with 5 to 12-µL of rAAV and allowed to adsorb to either a 300-mesh copper or 400-mesh carbon grid with Formvar and carbon coatings (Cat# FCF300-CU) for 3-min. The AAV was washed off by touching the grid, sample side down, to two drops of ddH_2_O (18-MΩ). 1-2% uranyl acetate in ddH_2_O (18-MΩ) was then dripped through a 0.2-µm mesh syringe filter over the grid allowing the third drop to remain on the grid for 1-min before most of the stain was wicked away with filter paper and allowed to dry. Grids were observed on either a JEOL JEM-1400 or JEOL JEM-1230 transmission electron microscope at 120-kV and photos were taken using Gatan Orius digital cameras (either 2K-x-2K or 4K-x-4K). Packaging percentage determinations from TEM images were measured in ImageJ v2.0.0-rc-43/1.52b^64^.

### False-colored structural capsid PTM mapping

The rAAV8 PTMs shown in Figure 1 were mapped onto the PDB AAV8 capsid structure 3RA8^17^ using Pymol v1.7.6.0. The rAAV1 PTMs shown in Figure S8 were mapped onto the PDB AAV1 capsid structure 3NG9^65^. Exterior capsid views have all chains represented, while internal cross-section views have chains surrounding a cylinder at the 5-fold symmetry axis removed exposing the interior capsid lumen with fogging cued to enable depth perception of the interior. Online UbPred software (http://www.ubpred.org) was used to predict the possible ubiquitination sites on AAV1 and AAV8 capsid proteins^20^. The output of the analysis is the prediction of the lysine residues important for ubiquitination within the indicated AAV serotype capsid sequence. There were 11 lysines on rAAV1 VP3: K258, K459, K491, K493, K508, K528, K533, K545, K567, K666 and K707. There were 8 lysines on rAAV8 VP3: K259, K333, K510, K530, K547, K569, K668 and K709.

### Cryo-EM data acquisition and image processing

Purified rAAV8 samples (human-produced empty and full, baculovirus-*Sf9* produced empty and full) were applied to Lacey carbon Quantifoil grids (EMS Cat# Q225CR-06) and flash frozen. Images were collected using an FEI Titan Krios G3i Cryo Transmission Electron Microscope operating at 300-kV with a Gatan K2 Summit detector and GIF using a 20-eV slit width. Movies were collected in counting mode with the automated imaging software EPU and a pixel size of 1.07 Å. 30 frames were collected in a 6-sec exposure with a per frame dose rate of 2.2 e^-^/Å^2^ for a total dose of ∼66 e^-^/Å^2^. A total of 1,647 movies were collected for human-produced rAAV8 and 396 movies were collected for baculovirus-*Sf9* produced rAAV8. Movies were drift-corrected using MotionCor2^66^. All image processing was performed using cisTEM^67^. No symmetry was applied for initial 2D classification and icosahedral symmetry was applied for 3D classification and the final reconstruction. The previous crystal structure determined for rAAV8 (PDB 3RA8) was used as an initial starting point for all models. Residues were manually fit in Coot^68^ and refined against the cryo-EM maps using Phenix.real_space_refine^69^. The 4 new structures have been deposited in both PDB and EMDB: human HEK293-produced full AAV8 capsid (PDB-6PWA; EMD-20502); human HEK293-produced empty AAV8 capsid (PDB-6U20; EMD-20615); baculovirus-*Sf9* produced full AAV8 capsid (PDB-6U2V; EMD-20626); baculovirus-*Sf9* produced empty AAV8 capsid (PDB-6UBM; EMD-20710).

### Signal peptide determination

Putative ER signal peptides of the AAV VP1 polypeptide were found using LocSig Database software (http://genome.unmc.edu/LocSigDB)^31^. Only signals within the first 100 amino acid residues of VP1 from the N-terminal end were assessed.

### Gene ontology analysis on host cell protein impurities

Cumulative HCP impurities from all vector lots analyzed by LC-MS/MS (human impurities observed in human vector preparations; human impurities observed in baculovirus-*Sf9* vector preparations; *Spodoptera frugiperda* impurities observed in baculovirus-*Sf9* vector preparations; baculoviral impurities observed in baculovirus-*Sf9* vector preparations) were ranked by PSM. For human impurities, those with a PSM >3 were analyzed by gene ontology functional enrichment analysis using g:Profiler^70^, version e94_eg41_p11_9f195a1, with g:SCS multiple testing correction method applying significance threshold of 0.05 and an ordered query to generate Figure S10a. Only impurities from *Homo sapiens* could be assessed with these bioinformatic tools as both *Spodoptera frugiperda* and *Autographa californica multiple nuclepolyhedrovirus* (NPVAC, baculovirus) were not supported organisms. Gene ontologies for *Sf9* and *NPVAC* were assessed manually using UniProtKB.

### Whole genome bisulfite sequencing of packaged rAAV genomes

Total packaged gDNA was extracted from 1E11 full rAAV particles from two productions of ssAAV8-EF1ɑ-FLuc-WPRE-hGHpA, produced from each manufacturing platform and purified from cell lysates. To first remove unincorporated DNA, ssAAV vector samples were incubated with 40-U exonuclease (Lucigen Cat#E3101K) and 10-U endonuclease (Lucigen Cat#DB0715K) for 30-min at 37°C and then incubated at 65°C for 10-min to stop the digestion. The entire 200-µL reaction was used as input for ssAAV gDNA extraction using a MinElute Virus Spin Kit (Qiagen Cat#57704). 50-ng of ssDNA from each sample were bisulfite converted using the EZ Methylation Gold kit (Zymo Cat#D5005) according to the manufacturer’s protocol. Whole genome bisulfite sequencing (WGBS) libraries were generated from the converted samples with the TruSeq DNA Methylation kit (Illumina Cat#EGMK81312). Consistent fragment size distribution of the resulting libraries was assessed via micro-capillary gel electrophoresis on a Bioanalyzer 2100 (Agilent Cat#G2939BA). Libraries were quantified by Qubit fluorometry (LifeTechnologies Cat#Q33240), and pooled in equimolar ratios. The library pool was quantified via qPCR with a KAPA Library-Quant kit (Kapa Biosystems Cat#KK4824). Each sample was split in half and half were sequenced on an Illumina HiSeq 4000 system for single-end 100-bp reads, and half were sequenced on an Illumina NovaSeq 6000 system for paired-end 2×150-bp reads. Raw reads were subjected to quality control. Adapters were removed using scythe version c128b19 (https://github.com/vsbuffalo/scythe). Bases with quality <30 were trimmed from the 3’-end of the reads using sickle version 85e5117 (https://github.com/najoshi/sickle). Reads <30bp in length after trimming were discarded. BS-Seeker2^71^ (version 2.1.5) was used in Bowtie2 mode to map all reads that passed quality control onto the AAV genome. All cytosine sites that were covered by reads were extracted using the BS-Seeker2 “bs_seeker2-call_methylation.py” program. Cytosine sites that were not covered by all samples were excluded from further analysis. Differential methylation at each cytosine site was tested using Fisher’s exact test. Multiple test correction was carried out using the Benjamini-Hochberg FDR^72^ approach. Coverage spanned 100% of the rAAV transfer vector genome reference sequence. Figure 3 was generated with circus^73^ (version 0.69-9).

### Mice

Adult SCID-Balb/cJ mice (CBySmn.CB17-Prkdcscid/J) of each sex were purchased from The Jackson Laboratories (Cat#1803). *Fah/Rag*2*/Il2rgc* (FRG) deficient mice of each sex on a C57BL/6J background transplanted with human hepatocytes^74^ of each sex were a generous gift from Yecuris (www.yecuris.com). All mice were housed and maintained in specific-pathogen-free barrier facilities. FRG mice were maintained on irradiated high-fat low-protein mouse chow (Envigo Teklad Global 16% Protein Cat#2916) *ad libitum* to decrease flux through the tyrosine pathway. Following human hepatocyte transplantation and prior to AAV administration, FRG mice were maintained on a rotating water schedule: 1-week of 3.25% dextrose (Sigma Cat#G6152) water, 1-week of 3.25% dextrose + 100mg/mL cotrimoxazole (Sigma Cat#A2487) water, and 1-week of 3.25% dextrose + 100mg/mL cotrimoxazole + 8mg/L Nitisinone (Yecuris Cat#20-0026) water. Following AAV administration and for the duration of the imaging studies, humanized FRG mice were maintained on 3.25% dextrose + 100mg/mL cotrimoxazole + 8mg/L Nitisinone water to ensure no liver turnover during the AAV transduction period. The Institutional Animal Care & Use Committees of UCSF and Stanford University approved all mouse procedures.

### Statistics

Statistical analyses were conducted with Prism v8.0.1 software. Experimental differences for Figures 4a-d and S1b were evaluated using a one-way ANOVA with Tukey’s correction for multiple comparisons assuming equal variance. Experimental differences for Figures 4e-h, 5, 6 and S12 were evaluated using a two-way ANOVA with Fisher’s LSD assuming equal variance. *P* values <0.05 were considered statistically significant. * *P* ≤ 0.05, ** *P* ≤ 0.01, *** *P* ≤ 0.001, **** *P* ≤ 0.0001.

