## Supplemental Figures and Small Tables for "Methods Matter -- Standard Production Platforms for Recombinant AAV Produce Chemically and Functionally Distinct Vectors"

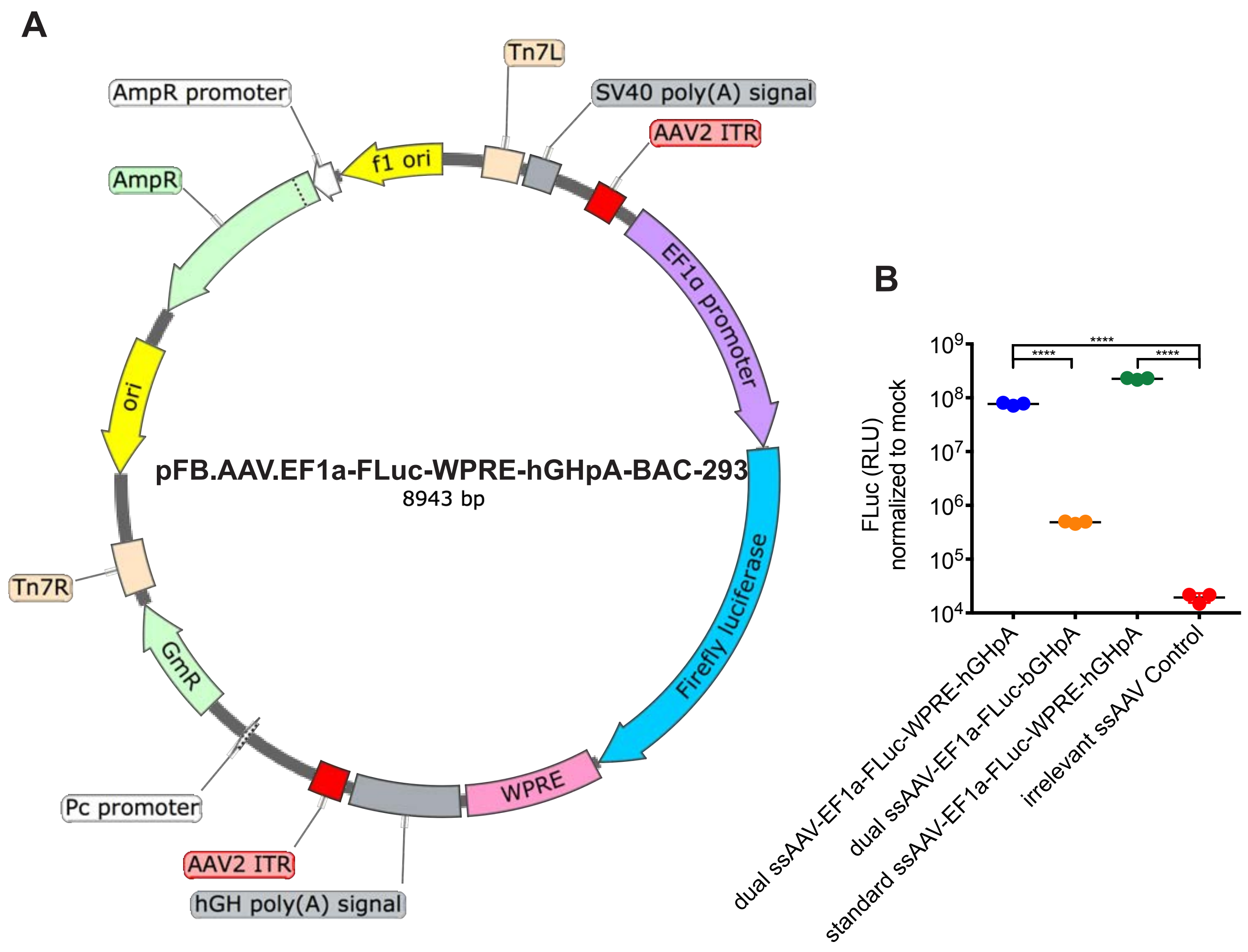


**Supplemental Figure 1. Custom dual-use plasmid pFB.AAV-EF1α-FLuc-WPRE-hGHpA-BAC-293. (A)** The plasmid was designed to have the necessary backbone components for use in both transient transfection into human HEK293 cells, as well as to produce live baculovirus. This ensured that the rAAV transfer vector sequence was identical between the two production platforms. Using this plasmid, each platform will produce ssAAV with an EF1α promoter driving expression of Firefly luciferase, followed by a WPRE to improve expression, and a standard hGHpA signal embedded between AAV2 ITRs. Within the backbone, two sets of important elements were included: AmpR expression components for growth in ampicillin-containing bacterial cultures for amplifying the plasmid itself; as well as gentamicin expression components and Tn7 transposition sites to allow for replication and transposition in insect cells during baculovirus production. The EF1α promoter was chosen as it is a strong ubiquitous non-tissue specific promoter that expresses well in a variety of cells/tissues both in vitro and in vivo to allow for maximal utility in the lab when making comparisons between species of interest and experimental conditions. Firefly luciferase was chosen as the reporter gene as it is easily detectable both in vitro and in vivo with highly sensitive commercial kits and common live bioluminescent imaging modalities in all tissues. WPRE inclusion enabled better expression detection, even in tissues/cells with traditionally low rAAV transduction frequencies. **(B)** FLuc expression data comparing the standard parental plasmid (only usable in the human production method) to the two dual test constructs with and without the WPRE demonstrating that the WPRE element is needed to achieve robust expression to enable comparative expression studies.

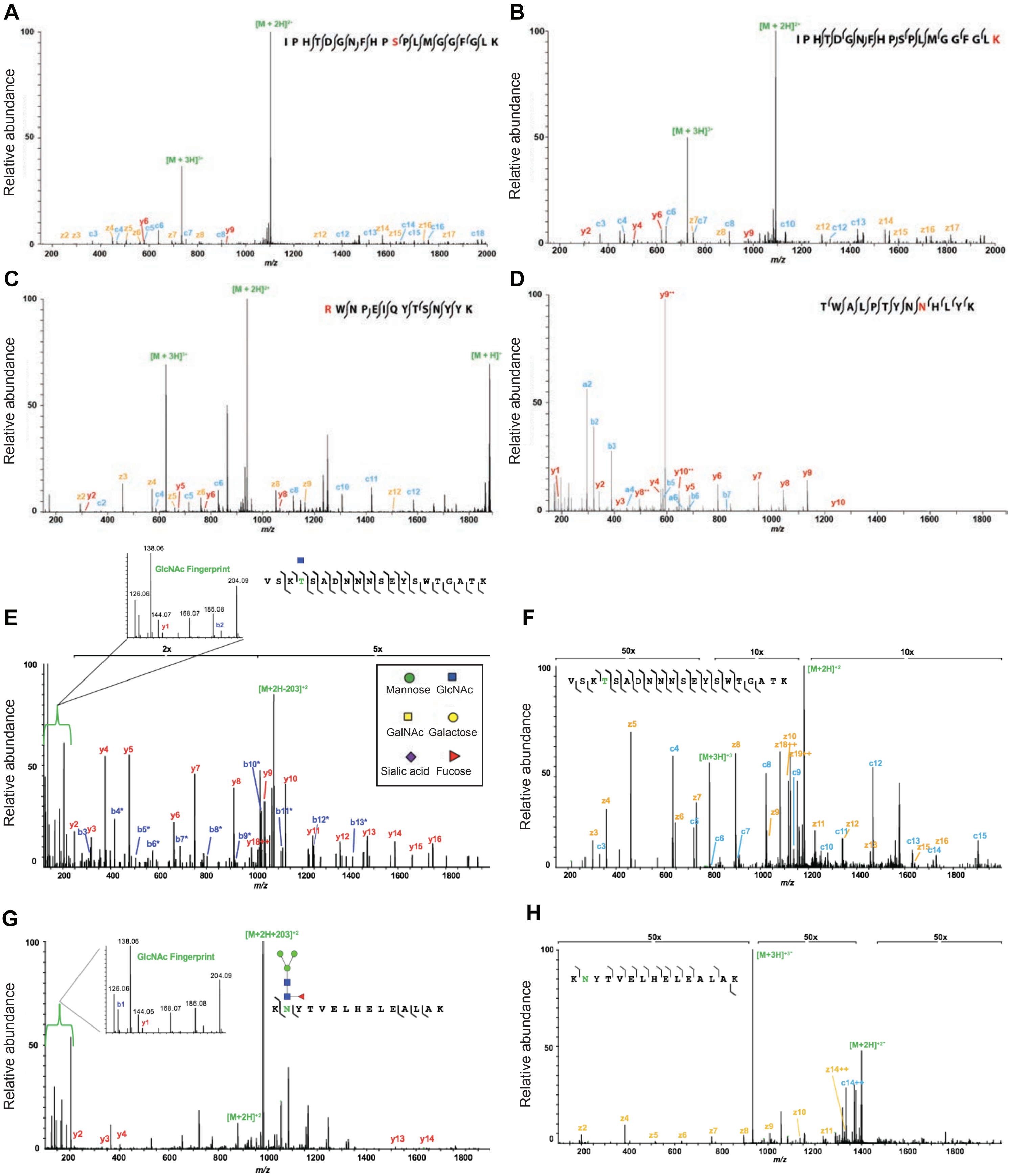


**Supplemental Figure 2: Example spectra for PTMs observed on rAAV capsids and HCP impurities.** **(A)** Example of a MS/MS fragmentation spectrum of a typical phosphorylated rAAV peptide sequence at the serine residue highlighted in red. **(B)** Example of a MS/MS fragmentation spectrum of a typical acetylated rAAV peptide sequence at the lysine residue highlighted in red. **(C)** Example of a MS/MS fragmentation spectrum of a typical methylated rAAV peptide sequence at the arginine residue highlighted in red. **(D)** Example of a MS/MS fragmentation spectrum of a typical deamidated rAAV peptide sequence at the asparagine residue highlighted in red. **(E)** Example of a typical HCD mass spectrum of a O-GlcNAcylated rAAV peptide at the initial threonine residue highlighted in green. Fragment ions that define the complete amino acid sequence are labeled as b and y. Those that have lost the O-GlcNAc moiety are labeled with an asterisk. **(F)** The associated ETD mass spectrum for (E). **(G)** Example of a typical HCD mass spectrum of an N-glycosylated *Sf9* HCP impurity peptide from ferritin with a N-GlcNAc2-Fuc-Man3 glycoform at the highlighted asparagine residue in green. Branching patterns were predicted based on known glycan structures along with sequential neutral losses from the mass spectra. **(H)** The associated ETD mass spectrum for (G).

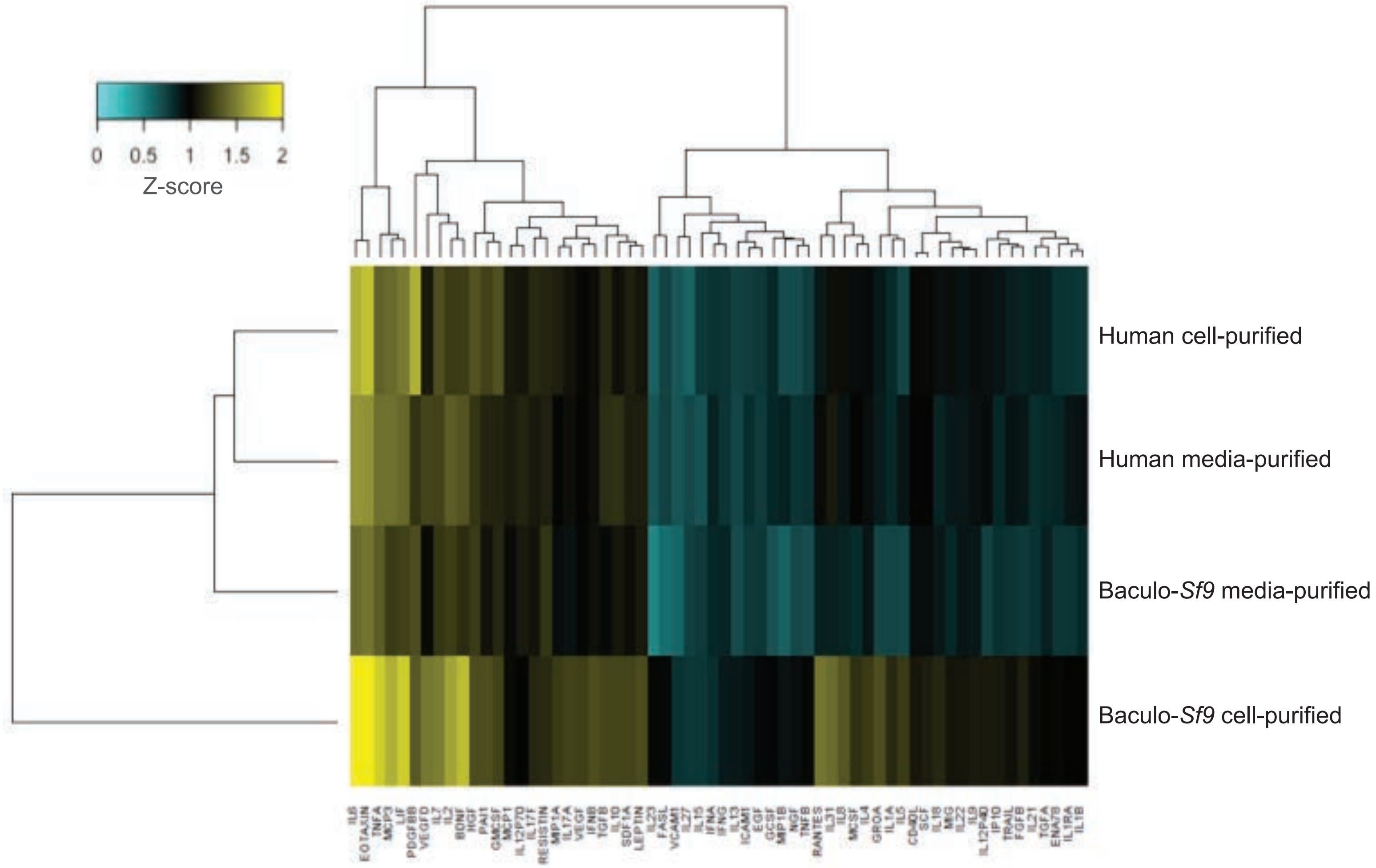


**Supplemental Figure 3: Cytokine/chemokine responses of primary human fibroblasts in response to rAAV transduction by different vector lot sources.** A heat map shows the results of a quantitative human 63-plex Luminex assay measured across four rAAV8 samples at 24 hours post-transduction. Heatmap cells are colored according to the normalized fold change Z score. The clustergram shows hierarchical clustering of virus lots based on similarity.

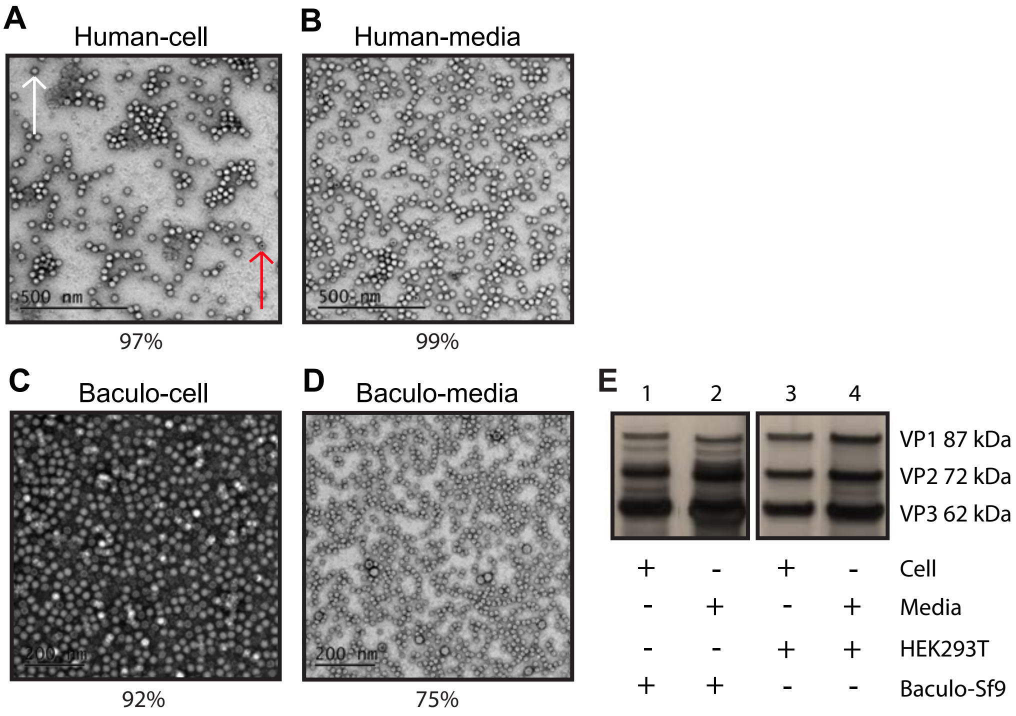


**Supplemental Figure 4: Comparative proteomic profiles of repeat rAAV8 vector lots manufactured with human and first-generation baculovirus-Sf9 production platforms.** **(A)** Negative staining and TEM imaging of human rAAV8 cell-purified vector (pair with B). White arrow denotes a full capsid, red arrow denotes an empty capsid, for reference for panels A-D (in all cases, the percent of full capsids containing a genome denoted below the representative TEM image above; Magnification is 20,000X). **(B)** Negative staining and TEM imaging of human rAAV8 media-purified vector (pair with A). **(C)** Negative staining and TEM imaging of baculo-*Sf9* rAAV8 cell-purified vector (pair with D). **(D)** Negative staining and TEM imaging of baculo-*Sf9* rAAV8 media-purified vector (pair with C). **(E)** Silver stain of capsid VP protein species present in vector lots from A-D. Denoted below whether capsids were purified from cell lysates or media supernatants, and which production platform was used for vector manufacturing. The truncated VP species with the baculo-*Sf9* vector are particularly prominent.

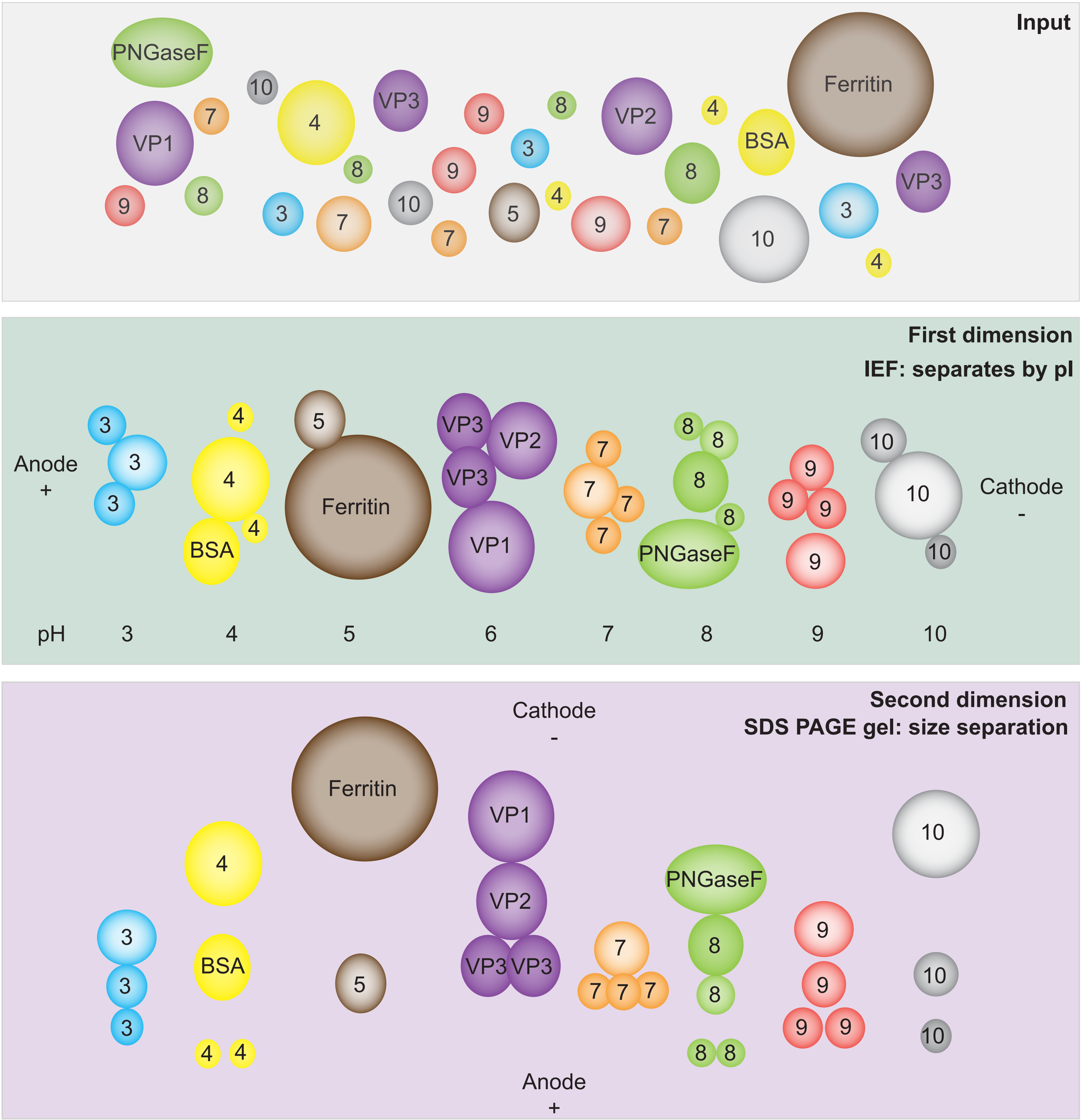


**Supplemental Figure 5: Schematic of 2D-gel electrophoresis process.** Vector lot samples composed of rAAV capsid proteins, host cell protein impurities and any process contaminants were loaded into precast pH gradient gel strips and run in the presence of current such that proteins migrate to the well where their isoelectric point (pI) matches the pH in the well. Once done migrating "in gel" in the first dimension, samples in each well are removed via pipet and loaded on their own lane on a standard SDS PAGE gel to separate by size in the second dimension. Running "in gel" rather than "in solution" creates bands rather than the more traditional dots.

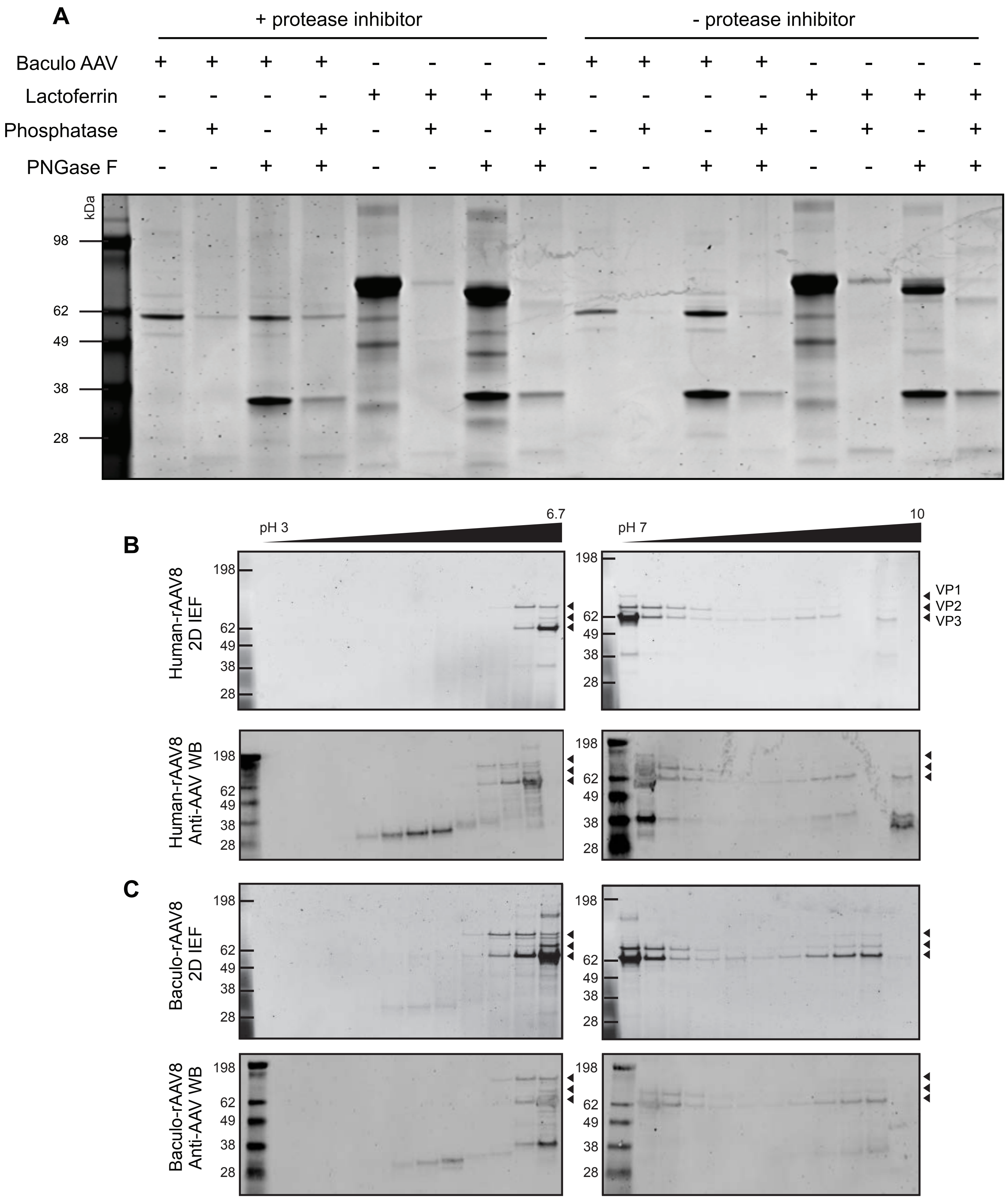


**Supplemental Figure 6: 2D-gel electrophoresis control +/- protease inhibitor to confirm the lack of host protease activity.** **(A)** An rAAV8 sample produced with the baculo-*Sf9* platform was subjected to standard in-gel 2D-gel electrophoresis with and without pretreatment with protease inhibitors to assess the potential for host protease action being responsible for banding patterns seen in Figure 1K,L. Experimental samples and controls: baculo-rAAV8 (VP1 87 kDa, VP2 72 kDa, VP3 62 kDa); lactoferrin (80 kDa); lambda protein phosphatase (25 kDa); PNGase F (34.8 kDa). **(B)** 2D gel images from human-produced rAAV8 from Figure 1K with corresponding anti-AAV VP1/VP2/VP3 western blot beneath. **(C)** 2D gel images from baculo-*Sf9* produced rAAV8 from Figure 1L with corresponding anti-AAV VP1/VP2/VP3 western blot beneath.

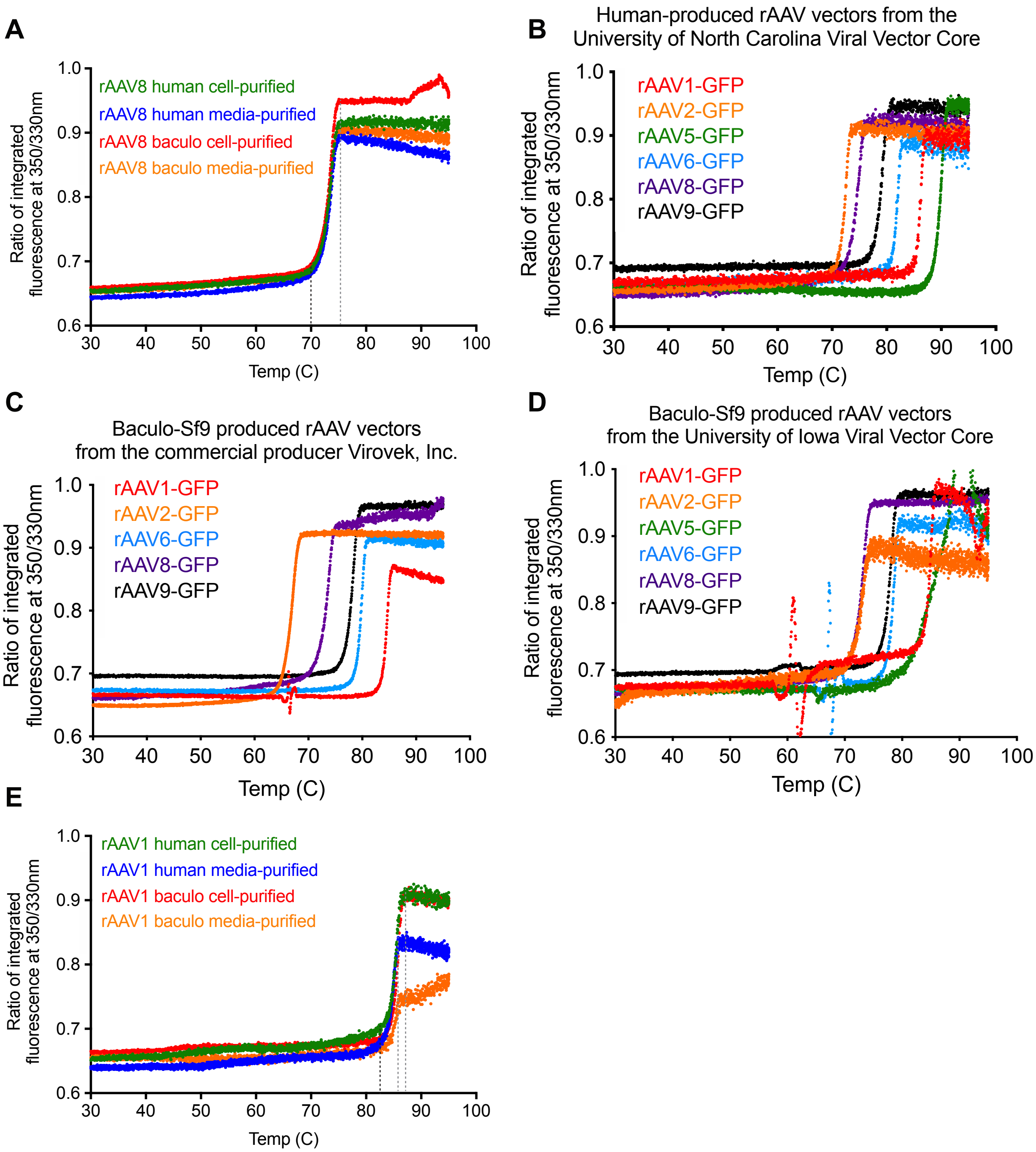


**Supplemental Figure 7: Thermal capsid melt curves to predict capsid stability. (A)** Full thermal capsid protein melt curves for rAAV8 vectors from 30-95C, rather than the cropped temperature curves shown in Figure 1M. Melt curve initiation is shown with a dashed black line, and final Tm is shown with a dashed grey line. No difference was seen for rAAV8 vectors produced in either manufacturing platform or purification source. **(B)** Thermal capsid melt curves for commonly used rAAV serotypes produced using transient transfection of human HEK293 cells at the UNC Viral Vector Core facility. All vectors contain the same CMV-GFP transfer cassette. The same legend color was used for panels B-D: rAAV1 (red), rAAV2 (orange), rAAV5 (green), rAAV6 (blue), rAAV8 (purple), and rAAV9 (black). **(C)** Thermal capsid melt curves for commonly used rAAV serotypes produced using live baculoviral infection of suspension *Sf9* cells at the commercial supplier Virovek, Inc. All vectors contain the same CMV-GFP transfer cassette. The aberration in signal from rAAV1 seen at 66C is common when capsids aggregate and/or when other protein contaminants are present. **(D)** Thermal capsid melt curves for commonly used rAAV serotypes produced using live baculoviral infection of suspension *Sf9* cells at the University of Iowa Viral Vector Core facility. All vectors contain the same CMV-GFP transfer cassette. Similar to the signal seen with Virovek baculoviral preparations, rAAV1 and rAAV6 had aberrant signal due to aggregation and/or contaminants. **(E)** Full thermal capsid protein melt curves for rAAV1 vectors from 30-95C, rather than the cropped temperature curves shown in Figure S8K. Melt curve initiation is shown with a dashed black line, and final Tm is shown with dashed grey lines. No difference was seen for rAAV1 vectors produced in either manufacturing platform.

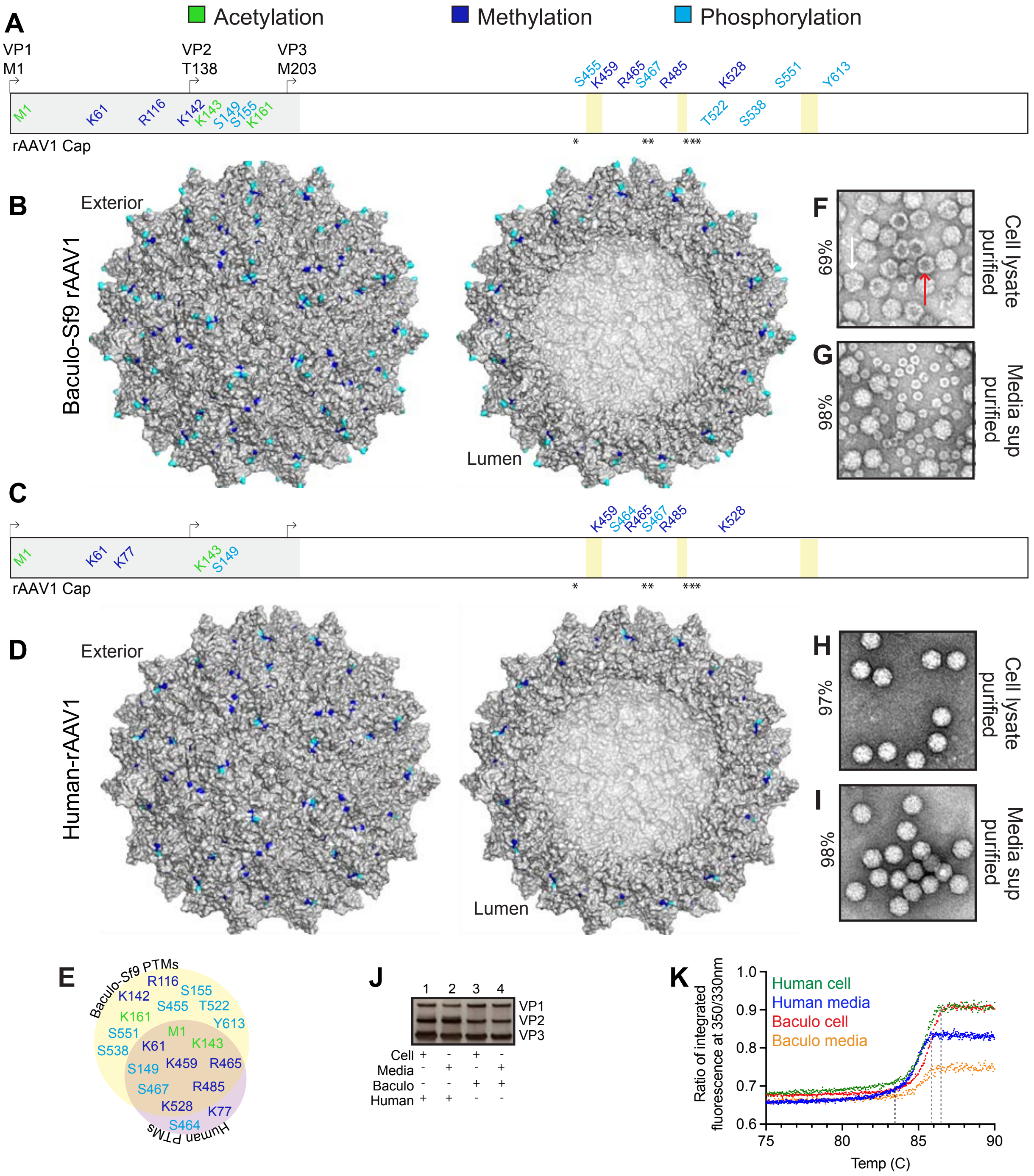


**Supplemental Figure 8: AAV1 vector preparations manufactured with the human and baculovirus-*Sf9* production platforms exhibit differential PTM profiles. (A)** PTM identities and residue positions along the length of the rAAV1 polypeptide from the N to C-terminus in baculo-*Sf9* vector. PTMs colored by type (acetylation = green, methylation = blue, phosphorylation = cyan). Residues above the sequence are externally facing on the capsid. Residues below are lumenal or buried. Residues within the grey box from residues 1-217 represent the disordered region of AAV1 yet to be crystallized. The six residues involved in sialic acid primary receptor binding (N447, S472, V473, N500, T502, and W503) are noted with an asterix. The three capsid motifs with known high antigenicity are denoted with yellow boxes: (456-AQNK-459), (492-TKTDNNNS-499), and (588-STD-PATGDVH-597). **(B)** Cumulative capsid PTMs observed from all baculo-*Sf9* rAAV1 lots, purified from both cell lysates and media. Color code as in (A). **(C)** Same as (A) but with human-produced rAAV1. **(D)** Same as (B) but with human rAAV1. **(E)** Shared and unique capsid PTMs for rAAV1 produced in the baculo-*Sf9* (yellow) and human systems (purple). Color code as in (A). **(F)** Negative staining and TEM imaging of baculo-*Sf9* rAAV1 cell-purified vector. White arrow denotes a full capsid, red arrow denotes an empty capsid, for reference for panels F-I (percent full capsids noted to the left). Magnification is 20,000X. **(G)** Same as (F) but media-purified vector. **(H)** Same as (F) but with human rAAV1 cell-purified vector. **(I)** Same as (H) but with media-purified vector. **(J)** Silver stain of capsid VP species present in vector lots from panels F-I. **(K)** Thermal capsid melt curves for rAAV1 vectors shown from 75-90C, full melt curves from 30-95C are in Figure S7E. Tm initiation = dashed black line; final Tm = dashed grey line.

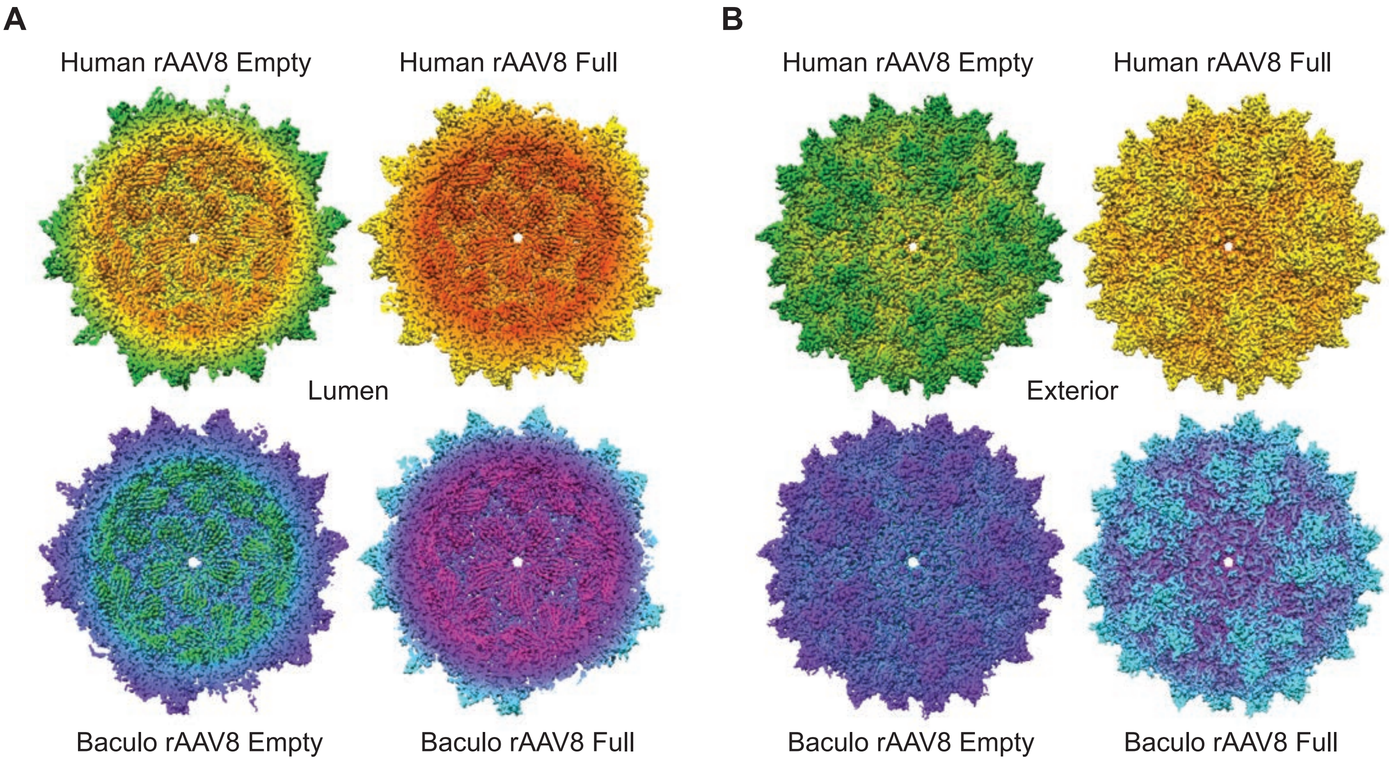


**Supplemental Figure 9: Additional cryo-EM structural data from the four new rAAV8 structures of full and empty capsids from baculovirus-*Sf9* and human-produced vectors. (A)** Lumenal cutaway capsid views centered on the interior base of the capsid cylinder pore of four new rAAV8 structures determined by cryo-EM. Vectors from each manufacturing platform, as well as empty and full capsids are shown. Human empty capsids are colored in orange, yellow, green; human full capsids are red, orange, yellow; baculo-*Sf9* empty capsids are green, blue, purple; and baculo-*Sf9* full capsids are pink, purple, cyan. **(B)** Exterior capsid views centered on the capsid cylinder pore of the four rAAV8 structures determined by cryo-EM. Same color scheme as in (A).

A

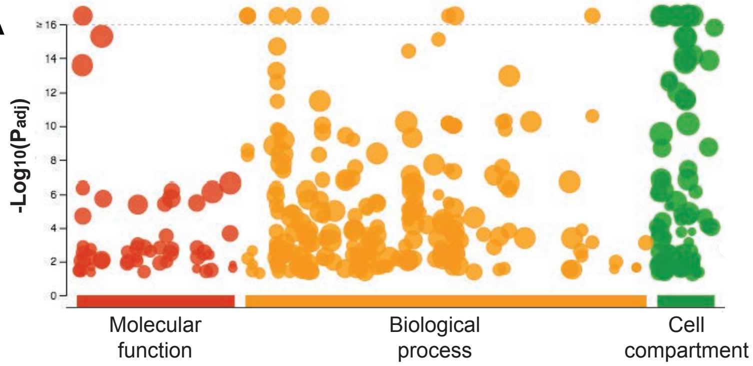


B

| | *H.sapiens* | *S.frugiperda* | *NPVAC* (Baculovirus) |
|---|---|---|---|
| Molecular function | Nucleic acid binding<br>Protein binding | Endopeptidase activity<br>Protein binding | Structural<br>Polyhedra liberation |
| Biological process | RNA processing | Proteolysis<br>Response to unfolded protein | Capsid formation<br>Viral escape |
| Cell compartment | Nucleus<br>Cytosol | Proteasome<br>Nucleus/cytosol | Viron |

**Supplemental Figure 10: Cumulative comparative HCP impurity analysis. (A)** Gene ontology analysis for human HCP impurities from human produced vector preparations. Only impurities from *Homo sapiens* could be assessed with these bioinformatic tools as both *Spodoptera frugiperda* and *Autographa californica multiple nuclepolyhedrovirus* (NPVAC, baculovirus) were not supported organisms. Gene ontologies for *Sf9* and NPVAC were assessed manually using UniProtKB. **(B)** Comparative gene ontology analysis for the most common HCP impurities from each host type: *H.sapiens*, *S.frugiperda* and NPVAC (baculovirus).

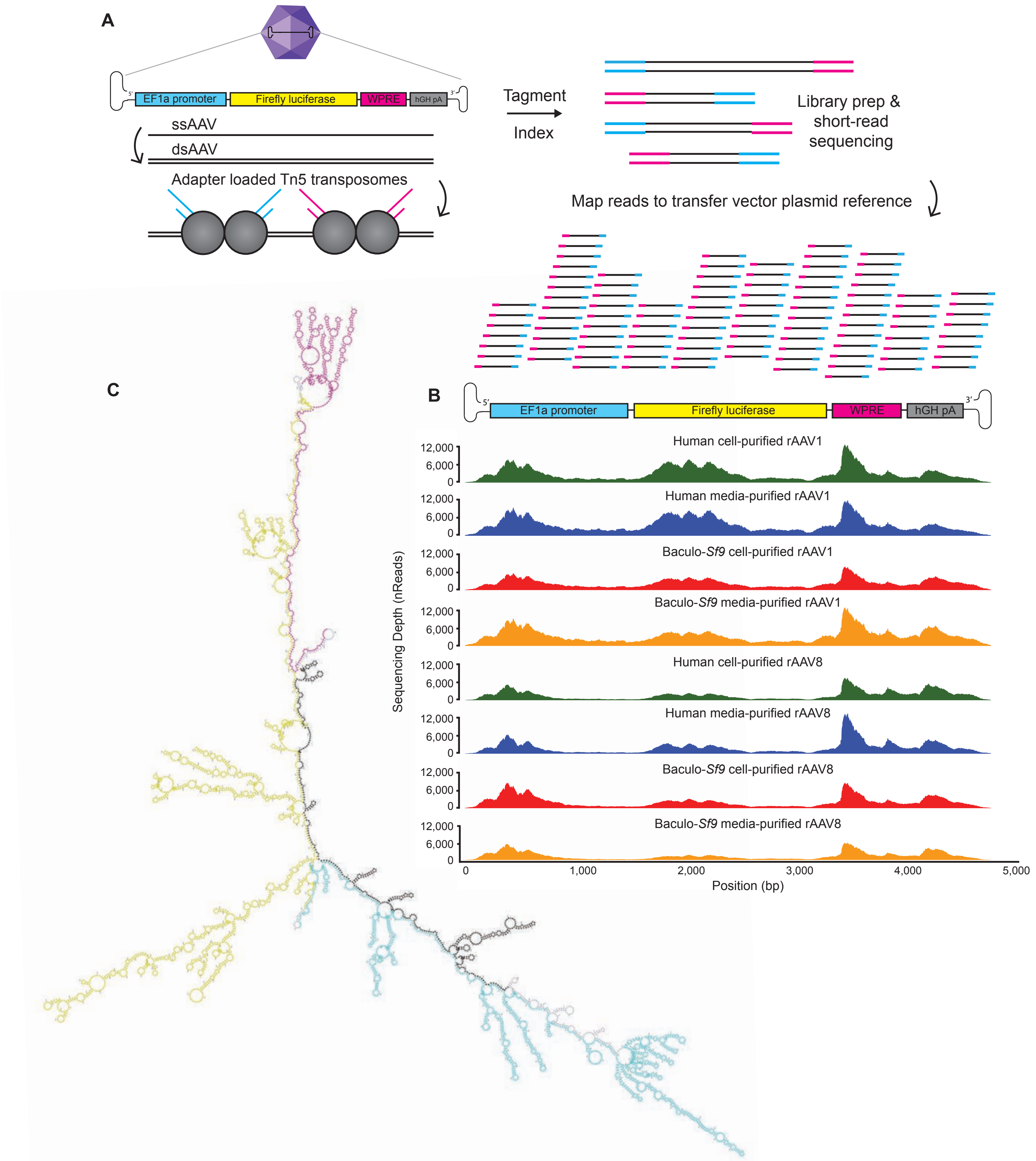


**Supplemental Figure 11: Packaged genome validation using Fast-Seq NGS method. (A)** Scheme of our new NGS validation protocol Fast-Seq used to validate whether packaged rAAV genomes, including ITRs, from human and baculo-produced vectors were full-length and identical to one another. **(B)** NGS sequence plots of depth (nReads) and coverage of packaged rAAV genomes, including ITRs. Sequencing validated that genomic payload sequence from human and baculo-*Sf9* produced vectors are identical and full-length. **(C)** Predicted ssDNA folding map of our packaged custom rAAV genome highlighting regions of complementarity and local secondary structure. Color coding matches the transfer vector in (B), wherein the EF1α promoter is cyan, the Firefly luciferase cDNA coding sequence is magenta, the WPRE is yellow, and the hGHpA element is black. Intervening sequences between these elements are shaded in lavender.

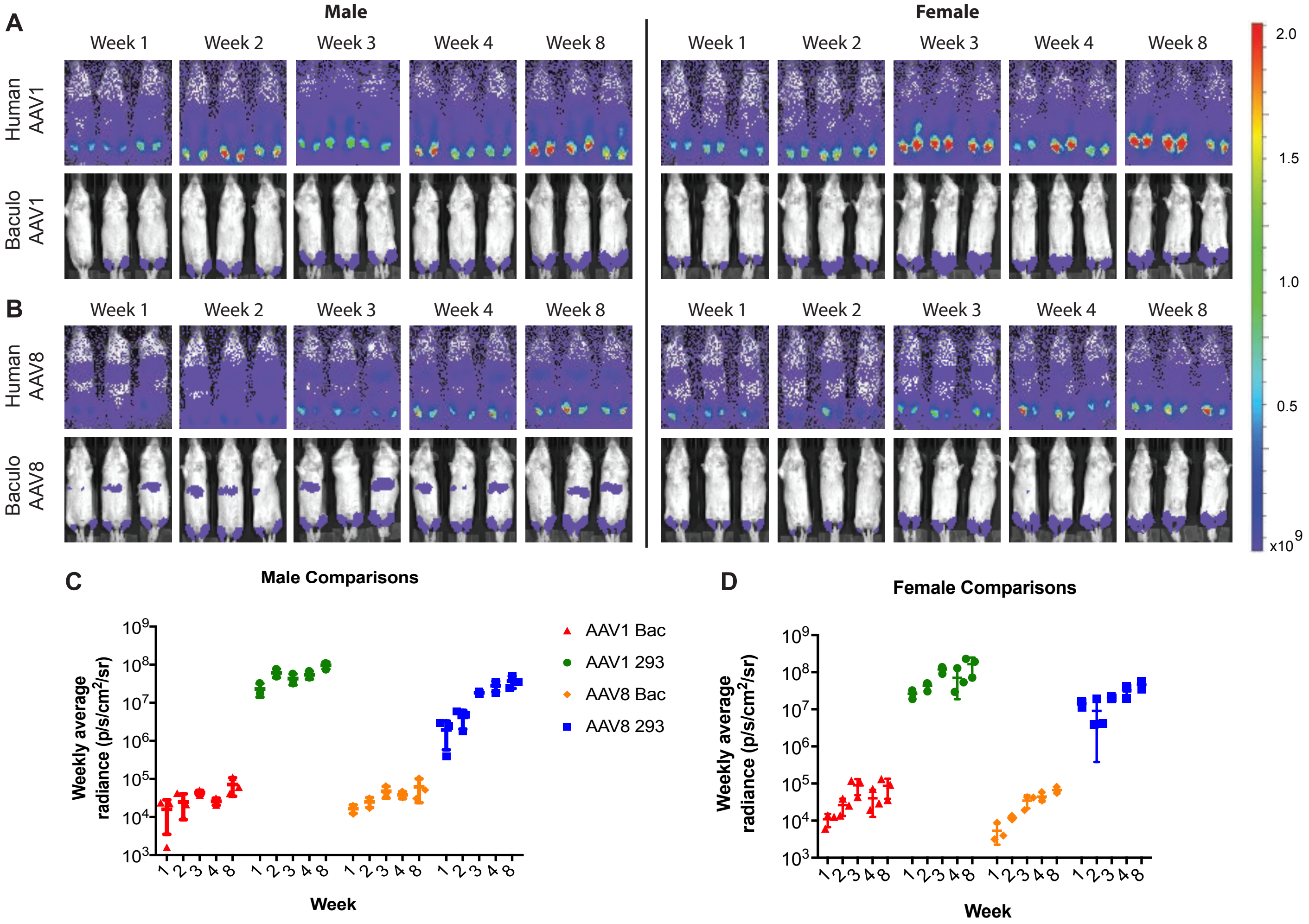


**Supplemental Figure 12: Additional *in vivo* data from Figure 4. (A)** Time course functional transduction FLuc expression from human and baculo-*Sf9* produced ssAAV1-EF1α-FLuc after IM administration (5E10 vg/mouse) in age-matched male and female sibling Balb/SCID mice. Mean radiance (p/s/cm2/sr) is displayed with all mice imaged on their ventral side on the same, shared scale. Image is shown highly blown out to enable seeing the low level of comparative expression in the baculo-treated mice (when compared to Figure 4E). **(B)** Time course functional transduction FLuc expression from human and baculo-*Sf9* produced ssAAV8-EF1α-FLuc after IM administration (5E10 vg/mouse) in age-matched male and female sibling Balb/SCID mice. Mean radiance (p/s/cm2/sr) is displayed with all mice imaged on their ventral side on the same, shared scale. Image is shown highly blown out to enable seeing the low level of comparative expression in the baculo-treated mice (when compared to Figure 4F). **(C)** Alternative representation to Figure 4G,H, now putting all serotypes on one graph and focusing on the differences in male mice. Each symbol represents the mean signal (+/- SD) from 3 mice. **(D)** Alternative representation to Figure 4G,H now putting all serotypes on one graph and focusing on the differences in female mice. Same color legend as (B). Each symbol represents the mean signal (+/- SD) from 3 mice.

### SUPPLEMENTAL TABLES

**Supplemental Table 1. All HCP impurities present in human and baculovirus-*Sf9* preparations of rAAV8.** All HCP impurities identified by LC-MS/MS for rAAV8 from both cell lysate and media-purified vector preparations. Excluded from the list are common process contaminants that occur in routine sample preparation (human keratin, trypsin, etc.), and impurities with mutations/aberrations such that they didn't appear in the main search library. Searches were conducted against the complete proteomes of *Homo sapiens*, *Spodoptera frugiperda*, *Autographa californica multiple nuclepolyhedrovirus* (baculovirus), *Adeno-associated virus* serotype 8, and common process contaminants (BSA from media, etc.). PSM = peptide spectral match.

*[Table is an attached excel file].*

| rAAV | Source | Produced in | Cell or media | Purification | Titer (vg/mL) | Lot # |
|---|---|---|---|---|---|---|
| AAV8-EF1α-FLuc | UIowa | HEK293 | Cell | I-MQ | 1.10E+13 | AAV3242 |
| | | | Media | S-TFF-I-MQ | 2.60E+13 | AAV3243 |
| | | Baculo-*Sf9* | Cell | I-MQ | 8.70E+13 | AAV3244 |
| | | | Media | S-TFF-I-MQ | 3.30E+12 | AAV3245 |

**Supplemental Table 2. Viral specifications for the rAAV8 vector lots used in Figure 1.** A single transfer vector plasmid lot (Figure S1) was used to produce vector in each system. Human rAAV vector lots were produced via the standard 3-plasmid transient transfection system in HEK293(FT) cells. Baculo-*Sf9* rAAV vector lots were produced via the second-generation baculovirus-infected *Sf9* cell system. All vector lots were produced with AAV2 *Rep* and AAV2 ITRs with single-stranded genome configurations. Abbreviations: I = iodixanol gradient; MQ = MustangQ column; TFF = tangential flow filtration; S = 100-kDa MWCO filter.

| rAAV | Source | Produced in | Cell or media | Purification | Titer (vg/mL) | Lot # |
|---|---|---|---|---|---|---|
| AAV8-EF1α-FLuc | UIowa | HEK293 | Cell | I-MQ | 2.80E+12 | AAV3098 |
| | | | Media | S-TFF-I-MQ | 6.00E+12 | AAV3099 |
| | | Baculo-*Sf9* | Cell | I-MQ | 3.50E+13 | AAV3101 |
| | | | Media | S-TFF-I-MQ | 1.00E+12 | AAV3103 |

**Supplemental Table 3. Viral specifications for the rAAV8 vector lots used in Figure S4.** A single transfer vector plasmid lot (Figure S1) was used to produce vector in each system. Human rAAV vector lots were produced via the standard 3-plasmid transient transfection system in HEK293(FT) cells. Baculo-*Sf9* rAAV vector lots were produced via the second-generation baculovirus-infected *Sf9* cell system. All vector lots were produced with AAV2 *Rep* and AAV2 ITRs with single-stranded genome configurations. Abbreviations: I = iodixanol gradient; MQ = MustangQ column; TFF = tangential flow filtration; S = 100-kDa MWCO filter.

| Source | rAAV | Produced in | Purification | Lot # |
|---|---|---|---|---|
| UNC Chapel Hill | AAV1-CBA-GFP | HEK293 | Column | AV6034 |
| | AAV2-CBA-GFP | | | AV6035 |
| | AAV5-CBA-GFP | | | AV6036 |
| | AAV6-CBA-GFP | | | AV6037d |
| | AAV8-CBA-GFP | | | AV6038 |
| | AAV9-CBA-GFP | | | AV6039 |
| Virovek | AAV1-CMV-GFP | Baculo-*Sf9* | CsCl | 15-621 |
| | AAV2-CMV-GFP | | | 16-017 |
| | AAV6-CMV-GFP | | | 13-195 |
| | AAV8-CMV-GFP | | | 14-365 |
| | AAV9-CMV-GFP | | | 15-409 |
| UIowa | AAV1-CMV-eGFP | Baculo-*Sf9* | I-MQ | AAV2820 |
| | AAV2-CMV-eGFP | | | AAV2601 |
| | AAV5-CMV-eGFP | | | AAV2844 |
| | AAV6-CMV-eGFP | | | AAV2588 |
| | AAV8-CMV-eGFP | | | AAV2846 |
| | AAV9-CMV-eGFP | | | AAV2646 |

**Supplemental Table 4. Biomanufacturing facilities used to determine the universality of rAAV capsid thermal stability in different capsid serotypes in Figure S7B-D.** Human rAAV vector lots manufactured at UNC were produced via transient plasmid transfection into suspension HEK293 cells in serum and antibiotic-free media. Insect rAAV vector lots manufactured at University of Iowa and Virovek were produced via live baculoviral infection of *Sf9* insect cells. All vector lots were produced with AAV2 *Rep* and AAV2 ITRs. Abbreviations: I = iodixanol gradient; MQ = MustangQ column; CsCl = cesium chloride density gradient; CBA = chicken b-actin promoter; CMV = cytomegalovirus promoter; GFP = green fluorescent protein.

**Supplemental Table 5. All HCP impurities present in human and baculovirus-*Sf9* preparations of rAAV1.** All HCP impurities identified by LC-MS/MS for rAAV1 from both cell lysate and media-purified vector preparations. Excluded from the list are common process contaminants that occur in routine sample preparation (human keratin, trypsin, etc.), and impurities with mutations/aberrations such that they didn't appear in the main search library. Searches were conducted against the complete proteomes of *Homo sapiens*, *Spodoptera frugiperda*, *Autographa californica multiple nucleopolyhedrovirus* (baculovirus), *Adeno-associated virus* serotype 1, and common process contaminants (BSA from the media, etc.). PSM = peptide spectral match.

*[Table is an attached excel file].*

| Production | Source | HCP impurity peptide | Unmodified peptide mass (Da) | Modified peptide mass (Da) | Hex NAc | Hex | Me-HexA | Fuc |
|---|---|---|---|---|---|---|---|---|
| Transient transfection in human HEK293 | Cell lysate | None | n/a | | | | | |
| | Media | None | n/a | | | | | |
| Live baculovirus infection of *Spodoptera frugiperda Sf9* insect cells | Cell lysate | K**N**YTVELHELEALAK (*Sf9* ferritin) | 1757.9432 | 3136.4374 | 2 | 6 | | |
| | | | | 3298.4838 | 2 | 7 | | |
| | | | | 2796.3262 | 2 | 3 | | 1 |
| | | | | 2634.2782 | 2 | 2 | | 1 |
| | | | | 1961.0278 | 1 | | | |
| | | AK**N**YTVELHELEALAK (*Sf9* ferritin) | 1828.9803 | 2867.3706 | 2 | 3 | | 1 |
| | | | | 2032.0702 | 1 | | | |
| | | **N**YTVELHELEALAK (*Sf9* ferritin) | 1629.8483 | 3332.43926 | 2 | 8 | | |
| | | | | 3170.388 | 2 | 7 | | |
| | | | | 3008.3616 | 2 | 6 | | |
| | | | | 1832.934 | 1 | | | |

**Supplemental Table 6. HCP impurities with N-linked glycans from human and baculo-*Sf9* produced preparations of rAAV1.** N-linked glycans (modified residue is bold) present on HCP impurities identified by LC-MS/MS for rAAV1 from both cell lysate and media-purified vector preparations. Excluded from the list are common process contaminants that occur in routine sample preparation (human keratin, trypsin, etc.), impurities with mutations/aberrations such that they didn't map to known proteins by BLASTp search, and any modification which could not be site-localized. HexNAc = N-acetylhexoseamine; Hex = hexose; Me-HexA = methylated hexuronic acid; Fuc = fucose.

| rAAV | Source | Produced in | Cell or media | Purification | Titer (vg/mL) | Lot # |
|---|---|---|---|---|---|---|
| AAV1-EF1α-FLuc | UIowa | HEK293 | Cell | I-MQ | 1.50E+12 | AAV3170 |
| | | | Media | S-TFF-I-MQ | 2.80E+12 | AAV3174 |
| | | Baculo-*Sf9* | Cell | I-MQ | 1.20E+13 | AAV3168 |
| | | | Media | S-TFF-I-MQ | 2.40E+12 | AAV3169 |

**Supplemental Table 7. Viral specifications for the rAAV1 vector lots used in Figure S8.** A single transfer vector plasmid lot (Figure S1) was used to produce vector in each system. Human rAAV vector lots were produced via the standard 3-plasmid transient transfection system in HEK293(FT) cells. Baculo-*Sf9* rAAV vector lots were produced via the baculovirus-infected *Sf9* cell system. All vector lots were produced with AAV2 *Rep* and AAV2 ITRs with single-stranded genome configurations. Abbreviations: I = iodixanol gradient; MQ = MustangQ column; TFF = tangential flow filtration; S = 100-kDa MWCO filter.

| rAAV8 sample type | # of micrographs | # of particles | Resolution (Å) |
|---|---|---|---|
| HEK293 Full | 1,645 | 63,471 | 3.3 |
| HEK293 Empty | 1,645 | 15,107 | 3.3 |
| Baculo-*Sf9* Full | 395 | 404 | 3.6 |
| Baculo-*Sf9* Empty | 395 | 2,359 | 3.3 |

**Supplemental Table 8. Summary of cryo-EM processing parameters for the four new rAAV8 structures.** Processing parameters for the four rAAV8 structures purified from cell lysates from each manufacturing platform, including both full (had a genome) and empty capsids (lacked an AAV genome). More detailed viral manufacturing specifications for the rAAV8 vector lots used are listed in Table S4. Å = angstrom.

| Source | rAAV | Produced in | Purification | Lot # |
|---|---|---|---|---|
| UIowa | AAV1-CMV-eGFP | HEK293 | I-MQ | AAV2762 |
| | AAV5-CMV-eGFP | | I-MQ | AAV2042 |
| | AAV6-CMV-eGFP | | CsCl | AAV2885 |
| | AAV8-CMV-eGFP | | I-MQ | AAV2337 |
| | AAV1-CMV-eGFP | Baculo-*Sf9* | I-MQ | AAV2820 |
| | AAV2-CMV-eGFP | | I-MQ | AAV2601 |
| | AAV4-CMV-eGFP | | I-MQ | AAV2771 |
| | AAV5-CMV-eGFP | | I-MQ | AAV2844 |
| | AAV6-CMV-eGFP | | I-MQ | AAV2588 |
| | AAV8-CMV-eGFP | | I-MQ | AAV2846 |

**Supplemental Table 9. Different rAAV capsid serotypes used to determine universality of capsid PTM deposition.** Human rAAV vector lots were produced with the classic 3-plasmid transient transfection system in HEK293(FT) cells. Insect rAAV vector lots were produced via the baculovirus-infected *Sf9* cell system. All vector lots were produced with AAV2 *Rep* and AAV2 ITRs, had single-stranded genome configurations, were purified only from cell lysate from a single facility at the University of Iowa Viral Vector Core. Abbreviations: I = iodixanol gradient; MQ = MustangQ column; CsCl = cesium chloride density gradient.

**Supplemental Table 10: PTMs present in human and baculovirus-*Sf9* preparations from various serotype vectors produced at a single facility in Table S9.** Manually validated PTMs identified by LC-MS/MS for serotypes listed in Table S9. Position numbering is for each serotype, unaligned to the other serotypes. Modification types: [+0.98492] = deamidation of Asn and Gln; [+14.01565] = methylation; [+15.99492] = oxidation of Met (often the result of processing, not a real PTM); [+42.01057] = acetylation; [+71.03711] = propionamido-Cys; [+79.96633] = phosphorylation.

*[Table is an attached Excel file].*

**Supplemental Table 11: HCP impurities present in human and baculo-*Sf9* preparations from various serotype vectors produced at a single facility in Table S9.** HCP impurities identified by LC-MS/MS for all serotypes listed in Table S9 from cell lysate purified vector preparations. Excluded from the list are common process contaminants that occur in routine sample preparation (human keratin, trypsin, etc.). Searches were conducted against the complete proteomes of *Homo sapiens*, *Spodoptera frugiperda*, *Autographa californica multiple nucleopolyhedrovirus* (baculovirus), adeno-associated virus serotypes 1/2/4/5/6/8, and common process contaminants (BSA from the media, etc.). PSM = peptide spectral match.

*[Table is an attached excel file].*

| Platform | AAV | Glycosylated HCP impurity peptide | Unmodified peptide mass (Da) | Modified peptide mass (Da) | Hex NAc | Hex | HexA | Fuc |
|---|---|---|---|---|---|---|---|---|
| Baculo-*Sf9* | AAV1 | K**N**YTVELHELEALAK (*Sf9* ferritin) | 1757.9436 | 1961.0236 | 1 | | | |
| | | **N**YTVELHELEALAK (*Sf9* ferritin) | 1629.8483 | 2668.2303 | 2 | 3 | | 1 |
| | | | | 1832.9298 | 1 | | | |
| | AAV5 | **N**YTVELHELEALAK (*Sf9* ferritin) | 1629.8483 | 2668.2303 | 2 | 3 | | 1 |
| | | | | 1978.9856 | 1 | | | 1 |
| | | | | 1832.9298 | 1 | | | |
| | | K**N**YTVELHELEALAK (*Sf9* ferritin) | 1757.9436 | 1961.0236 | 1 | | | |
| | | K**N**YTVELHELEALAK (*Sf9* ferritin) | 1757.9436 | 1961.0236 | 1 | | | |
| | AAV8 | K**N**YTVELHELEALAK (*Sf9* ferritin) | 1757.9436 | 1961.0236 | 1 | | | |
| Human HEK293 | AAV6 | VQPF**N**VTQGK (human lysosomal membrane glycoprotein-2) | 1117.6000 | 1320.6793 | 1 | | | |
| | | GHTLTL**N**FTR (human lysosomal membrane glycoprotein-1) | 1159.6218 | 1362.7011 | 1 | | | |
| | | AGHTLTL**N**FTR (human lysosomal membrane glycoprotein-1) | 1230.6589 | 1433.7382 | 1 | | | |
| | | AAIPSALDT**N**SSK (human galectin-3-binding protein precursor) | 1274.6587 | 1477.738 | 1 | | | |
| | | | | 1623.679 | 1 | | | 1 |
| | | T**N**STFVQALVEHVK (human prosaposin) | 1572.838 | 3314.4302 | 3 | 5 | 1 | 1 |
| | | | | 2611.2132 | 2 | 3 | | 1 |
| | | | | 2449.1604 | 2 | 2 | | 1 |
| | | | | 1921.9764 | 1 | | | 1 |

**Supplemental Table 12. Glycosylated HCP impurities from human and baculovirus-*Sf9* produced preparations of multiple AAV serotypes produced at one facility.** N-linked glycans (modified residue is bold) present on HCP impurities identified by LC-MS/MS for numerous common serotypes. Excluded from the list are common process contaminants that occur in routine sample preparation (human keratin, trypsin, etc.), impurities with mutations/aberrations such that they didn't map to known proteins by BLASTp search, and any modification that could not be site-localized. HexNAc = N-acetylhexoseamine; Hex = hexose; HexA = hexuronic acid; Fuc = fucose.

| rAAV | Source | Produced in | Purification | Lot # |
|---|---|---|---|---|
| AAV1-CMV-FLuc | UPenn | HEK293 | TFF-I-MQ | V2682TI |
| AAV1-CMV-rhCGB | SignaGen Labs | HEK293 | CsCl | AAV61882 |
| AAV2-CBA-GFP | UNC Chapel Hill | HEK293 | CsCl | AV6108 |
| AAV8-CBA-GFP | UNC Chapel Hill | HEK293 | CsCl | AV4909 |
| AAV8-CAG-TdTom | Stanford | HEK293T | CsCl | NPL1 |
| AAV8-CAG-TdTom | Stanford | HEK293T | CsCl | NPL2 |
| AAV8-CBA-GFP | ATCC reference strain | HEK293 | CsCl | 03112010SP2pcg |
| AAV8-CMV-FLuc | UPenn | HEK293 | TFF-I-MQ | V5489L |
| AAV2-CMV-GFP | UIowa | Baculo-*Sf9* | TFF-I-MQ | AAV2601 |
| AAV2-CMV-hAADC | UMass | Baculo-*Sf9* | AVB Sepharose | n/a |
| AAV8-CMV-GFP | UIowa | Baculo-*Sf9* | TFF-I-MQ | AAV2846 |
| AAV8-CMV-GFP | Virovek | Baculo-*Sf9* | CsCl | 15-129 |
| AAV8-CMV-FLuc | Virovek | Baculo-*Sf9* | CsCl | 16-255 |

**Supplemental Table 13. List of biomanufacturing facilities used to determine the universality of rAAV capsid PTM deposition regardless of manufacturer or method.** Human rAAV vector lots were produced via standard transient plasmid transfection into HEK293 cells. Insect rAAV vector lots were produced via live baculoviral infection of *Sf9* insect cells. All vector lots were produced with AAV2 *Rep* and AAV2 ITRs with single-stranded genome configurations. Note: ATCC lot was originally sourced from the University of Florida Powell Gene Therapy Center. Abbreviations: I = iodixanol gradient; MQ = MustangQ column; CsCl = cesium chloride density gradient; TFF = tangential flow filtration; rhCGB = rhesus chorionic gonadotropin; CBA = chicken b-actin promoter; TdTom = Td-Tomato; FLuc = firefly luciferase; CMV = cytomegalovirus promoter; hAADC = human aromatic L-amino acid decarboxylase.

| Made In | rAAV | Source | Peptide | Position |
|---|---|---|---|---|
| Human HEK293 | AAV8-GFP | ATCC | K.KRPVEP**S**[+79.96633]PQRSPDSSTGIGK.K | S149 |
| | | | K.KRPVEPSPQR**S**[+79.96633]PDSSTGIGK.K | S153 |
| | AAV1-rhCGB | SignaGen | K.**K**[+42.01057]RPVEQSPQEPDSSSGIGK.T | K143 |
| | | | K.KRPVEQ**S**[+79.96633]PQEPDSSSGIGK.T | S149 |
| | | | R.TQNQSGSAQN**K**[+14.01565]DLLFSRG.S | K459 |
| | | | R.TQNQSGSAQNKDLLFS**R**[+14.01565]G.S | K465 |
| | AAV2-GFP | UNC | K.KRPVEH**S**[+79.96633]PVEPDSSSGTGK.A | S149 |
| | AAV8-TdTom | Stanford | R.TQTTGGTAN**T**[+79.96633]QTLGFSQGGPNTMANQAK.N | T460 |
| | | | R.FFPSNGILIFG**K**[+14.01565]Q.N | K547 |
| | | | K.NTPVPADPP**T**[+79.96633]TFNQSK.L | T662 |
| | AAV8-TdTom | Stanford | **M**.[+42.01057]AADGYLPDWLEDNLSEGIR.E | M1 |
| | | | **M**.[+42.01057]AADGYLPDWLEDNLSEGIREW.W | M1 |
| | | | K.KRPVEP**S**[+79.96633]PQR.S | S149 |
| | | | K.RPVEPSPQRSPDS**S**[+79.96633]TGIGK.K | S157 |
| | AAV1-FLuc | U-Penn | **M**.[+42.01057]AADGYLPDWLEDNLSEGIREWWDLKPGAPKPKA.N | M1 |
| | | | K.GEPVNAADAAALEHD**K**[+14.01565]AYDQQLKAG.D | K77 |
| | | | K.**K**[+42.01057]RPVEQSPQEPDSSSGIGK.T | K143 |
| | | | K.RPVEQ**S**[+79.96633]PQEPDSSSGIGK.T | S149 |
| | | | M.ASH**K**[+42.01057]DDEDKFFPMSGVMIFGK.E | K528 |
| | AAV8-FLuc | U-Penn | **M**.[+42.01057]AADGYLPDWLEDNLSEGIREWWALKPGAPKPK.A | M1 |
| | | | K.KRPVEP**S**[+79.96633]PQR.S | S149 |
| | | | R.NSLANPGIA**M**[+15.99492]ATH**K**[+42.01057]DDEERFFPSNGILIFGK.Q | K530 |
| | | | M.ATH**K**[+42.01057]DDEERFFPS**N**[+0.98402]GILIFGK.Q | K530/N540 |
| | | | K.TTNPVATEEYGIVADNLQQQNTAPQIGTVN**S**[+79.96633]QGALPGMVWQNR.D | S600 |
| | | | Q.NRDV**Y**[+79.96633]LQGPIWAKIPHTDGNFHPSPLMGGFGLK.H | Y615 |
| Baculo-*Sf9* | AAV8-GFP | Virovek | R.FFPS**N**[+0.98402]GILIFG**K**[+14.01565]Q.N | N540/K547 |
| | | | R.DVYLQGPIWA**K**[+14.01565].I | K623 |
| | AAV8-FLuc | Virovek | K.KRPVEP**S**[+79.96633]PQRSPDSS**T**[+79.96633]GIGK.K | S149/T158 |
| | | | K.RPVEPSPQR**S**[+79.96633]PDSSTGIGK.K | S153 |
| | | | K.YHLNGR**N**[+0.98402]**S**[+79.96633]LANPGIAMATHKDDEER.F | N517/S518 |
| | | | R.FFPSNGILIFG**K**[+14.01565]Q.N | K547 |
| | | | L.KHPPPQILI**K**[+14.01565]NTPVPADPPTTFNQSK.L | K652 |
| | AAV2-hAADC | U-Mass | LQF**S**[+661.6639]QAGASDIRDQSR | S463 |
| | | | VSK**T**[+788.7003]SADNNNSEYSWTGATK | T491 |
| | | | LV(dN)PGPAMA**S**[+673.14]HKDDEEKFFPQSGVLIFGK | S525 |
| | AAV2-GFP | U-Iowa | K.KRPVEH**S**[+79.96633]PVEPDSSSGTGK.A | S149 |
| | AAV8-GFP | U-Iowa | R.NSLANPGIA**M**[+15.99492]ATH**K**[+42.01057].D | K530 |
| | | | R.FFPSNGILIFG**K**[+42.01057]QNAAR.D | K547 |
| | | | R.DNADYSDVMLT**S**[+79.96633]EEEIK.T | S564 |

**Supplemental Table 14: PTMs present in human and baculovirus-*Sf9* preparations from vectors listed in Table S13.** Manually validated PTMs identified by LC-MS/MS for vectors listed in Table S13 from both cell lysate and media-purified vector preparations. Position numbering is for each serotype, unaligned to the other serotypes. The bold amino acid preceding the bracket were modified with: [+0.98492] = deamidation of Asn and Gln; [+14.01565] = methylation; [+15.99492] = oxidation of Met (often the result of processing, not a real PTM); [+42.01057] = acetylation; [+79.96633] = phosphorylation; [+661.6639/673.14/788.7003] = O-linked (Ser/Thr) N-acetylhexoseamine.

**Supplemental Table 15: HCP impurities present in human and baculovirus-*Sf9* preparations from vectors listed in Table S13.** HCP impurities identified by LC-MS/MS for all serotypes and vectors listed in Table S13 from both cell lysate and media-purified vector preparations. Excluded from the list are common process contaminants that occur in routine sample preparation (human keratin, trypsin, etc.). Searches were conducted against the complete proteomes of *Homo sapiens*, *Spodoptera frugiperda*, *Autographa californica multiple nucleopolyhedrovirus* (baculovirus), *Adeno-associated virus* serotypes 1/2/8, and common process contaminants (BSA from the media, etc.). PSM = peptide spectral match.

*[Table is an attached excel file].*

**Supplemental Table 16: Cumulative human HCP impurities present in all human-produced rAAV vector preparations tested to date.** Cumulative combined list of all human HCP impurities identified by LC-MS/MS for all human-produced vectors from both cell lysate and media-purified vector preparations. Excluded from the list are common process contaminants that occur in routine sample preparation (human keratin, trypsin, etc.). PSM = peptide spectral match.

*[Table is an attached excel file].*

**Supplemental Table 17: Cumulative *Sf9* HCP impurities present in all baculo-*Sf9* produced rAAV vector preparations tested to date.** Cumulative combined list of all *Sf9* HCP impurities identified by LC-MS/MS for all baculo-*Sf9* produced vectors from both cell lysate and media-purified vector preparations. Excluded from the list are common process contaminants that occur in routine sample preparation (human keratin, trypsin, etc.). PSM = peptide spectral match.

*[Table is an attached excel file].*

**Supplemental Table 18: Cumulative baculoviral HCP impurities present in all baculo-*Sf9* produced rAAV vector preparations tested to date.** Cumulative combined list of all baculoviral HCP impurities identified by LC-MS/MS for all baculo-*Sf9* produced vectors from both cell lysate and media-purified vector preparations. Excluded from the list are common process contaminants that occur in routine sample preparation (human keratin, trypsin, etc.). PSM = peptide spectral match.

*[Table is an attached excel file].*

| Serotype | ER signal peptide | Position |
|---|---|---|
| AAV1 | Dx(1)E<br>[HK]x(1)K | 96-99<br>30-33, 74-77 |
| AAV2 | | 96-99<br>23-26, 74-77 |
| AAV3b | | 96-99<br>74-77 |
| AAV4 | | 95-98<br>29-32, 73-76 |
| AAV5 | | 95-98<br>29-32 |
| AAV6 | | 96-99<br>30-33, 74-77 |
| AAV8 | | 96-99<br>30-33, 74-77 |
| AAV9 | | 96-99<br>74-77 |
| AAV12 | | 96-99<br>74-77 |

**Supplemental Table 19. Putative ER signal peptides are conserved in all natural rAAV VP1 polypeptides.** Putative ER signaling peptides were assessed for their presence on the N-terminal VP1 sequence of known natural AAV serotypes. Dx(1)E refers to Asp-x-Glu where ‘x’ represents any amino acid, and (1) refers to the number of times the preceding sequence can occur. HK]x(1)K refers to His-Lys-x-Lys where ‘x’ represents any amino acid, and (1) refers to the number of times the preceding sequence can occur.

| Index 1 | Index 2 |
|---|---|
| TCAGTGGCGAGT | AAAGTATGCTAC |
| CCGCCGCAAGAA | AAAGGGAGCGTG |
| CTCGGGACTGTT | TAAGCAAAGCAA |
| TAGGGATGTAAT | ATAGATAGTTCC |
| GAAATCAAGCGG | TCTATGACCATC |
| TTTGGGTTGAAT | CAAATACACTGA |
| CGGGTCCTAAAC | ATTCGGATTACA |
| TTTCAGGCACTC | CTAAAGGTCAAA |

**Supplemental Table 20: Indices used for NGS sequencing of packaged rAAV genomes.** Sequencing indices placed within the i5 and i7 adapters to enable dual-indexed pooled Illumina sequencing via Fast-Seq.

| Metric | BM1 | HM1 | HC8 | BM8 | BC1 | HC1 | BC8 | HM8 |
|---|---|---|---|---|---|---|---|---|
| Genome size (bp) | 4,364 | 4,364 | 4,364 | 4,364 | 4,364 | 4,364 | 4,364 | 4,364 |
| Mean coverage | 2,851 | 3,080 | 1,395 | 1,145 | 1,936 | 2,545 | 1,734 | 1,659 |
| Median coverage | 2,587 | 2,524 | 1,238 | 930 | 1,661 | 1,973 | 1,408 | 1,380 |
| Genome coverage range, not including the ITRs | 849 - 11,052 | 850 - 10,194 | 164 - 6,410 | 160 - 4,935 | 578 - 6,701 | 540 - 10,869 | 324 - 7,321 | 230 - 11,479 |
| Genome coverage range of the ITRs | 190 - 2,110 | 135 - 1,778 | 38 - 1,193 | 28 - 1,112 | 96 - 1,451 | 74 - 1,411 | 81 - 1,641 | 51 - 989 |
| Number of SNPs | 0 | 0 | 0 | 0 | 0 | 0 | 0 | 0 |
| Number of MNPs | 0 | 0 | 0 | 0 | 0 | 0 | 0 | 0 |
| Number of indels | 0 | 0 | 0 | 0 | 0 | 0 | 0 | 0 |
| Number of others (SVs) | 0 | 0 | 0 | 0 | 0 | 0 | 0 | 0 |
| Number of multiallelic sites | 0 | 0 | 0 | 0 | 0 | 0 | 0 | 0 |
| Number of multiallelic SNP sites | 0 | 0 | 0 | 0 | 0 | 0 | 0 | 0 |

**Supplemental Table 21: Comparative metrics for NGS sequencing of packaged rAAV genomes by Fast-Seq including ITRs.** Metrics reported from Fast-Seq demonstrating no difference in the packaged genomes between vectors or serotypes packaged in either manufacturing platform. BM = baculo-*Sf9* media-purified; HM = human media-purified; HC = human cell-purified; BC = baculo-*Sf9* cell-purified; SV = structural variant; SNP = single nucleotide polymorphism; MNP = multi nucleotide polymorphism.

| rAAV | Source | Produced in | Cell or media | Purification | Titer (vg/mL) | Lot # |
|---|---|---|---|---|---|---|
| AAV1-CMV-FLuc-SV40pA | UPenn | HEK293 | Media | TFF-I-MQ | 7.06E+12 | V2682TI |
| | Virovek | Baculo-*Sf9* | Cell | CsCl | 2.28+13 | 15-093 |
| AAV8-CMV-FLuc-SV40pA | UPenn | HEK293 | Media | TFF-I-MQ | 7.89E+13 | V5489L |
| | Virovek | Baculo-*Sf9* | Cell | CsCl | 2.37E+13 | 16-255 |

**Supplemental Table 22. Viral specifications for the rAAV1 and rAAV8 vector lots used in Figure 4E-H.** A ssCMV-FLuc-SV40pA transfer vector was used at each facility. Human rAAV vector lots were produced via the standard 3-plasmid transient transfection system in HEK293 cells. Baculo rAAV vector lots were produced via the baculovirus-infected *Sf9* cell system. All vector lots were produced with AAV2 *Rep* and AAV2 ITRs with single-stranded genome configurations. Abbreviations: I = iodixanol gradient; MQ = MustangQ column; TFF = tangential flow filtration; CsCl = cesium chloride density gradient.

**Supplemental Table 23: Statistics from Figure 5.** Quantification of rAAV1 FLuc radiance in male and female mice from Figure 5. Each symbol = mean signal (+/- SD) from 3 mice. * $P \leq 0.05$, ** $P \leq 0.01$, *** $P \leq 0.001$, **** $P \leq 0.0001$.

*[Table is an attached excel file].*

**Supplemental Table 24: Transplantation details for humanized liver mice.** Information from the humanized liver mice manufacturer Yecuris Inc on key parameters for the human hepatocyte donors and recipient mice. FRG = *Fah/Rag2/Il2rgc* deficient mice; HHF = healthy human female; HHM = healthy human male.

*[Table is an attached excel file].*

**Supplemental Table 25: Database of all approved advanced biologic therapies that contain proteins in their formulation.** Detailed list of the nearly 400 approved protein-containing advanced therapy medicinal products for use in humans (database was updated as of date of publication in May 2020). Includes vaccines, antibodies, antibody-drug conjugates, immune globulins, anti-venoms and anti-toxins, gene and cell therapies, hormones, blood products and enzymes separated into different tabs based on therapy type. Listed for each therapeutic are the name (including trade names and proper names), cell line and species of production, manufacturer, approval year, target/indication, and in some cases the route of administration.

*[Table is an attached excel file].*
